# Multicellular Proportional-Integral-Derivative Control for Robust Regulation of Biological Processes

**DOI:** 10.1101/2024.10.25.620164

**Authors:** Vittoria Martinelli, Davide Fiore, Davide Salzano, Mario di Bernardo

**Affiliations:** Department of Electrical Engineering and Information Technology University of Naples Federico II Via Claudio 21 80125 Naples Italy; Department of Mathematics and Applications “R. Caccioppoli” University of Naples Federico II Via Cintia Monte S.Angelo 80126 Naples Italy; SSM - School for Advanced Studies Via Mezzocannone 4 80138 Naples Italy

**Keywords:** Synthetic biology, biological control, genetic regulatory systems, PID control

## Abstract

This paper presents the first implementation of a Proportional-Integral-Derivative (PID) biomolecular controller within a consortium of different cell populations, aimed at robust regulation of biological processes. By leveraging the modularity and cooperative dynamics of multiple engineered cell populations, we develop a comprehensive in silico analysis of the performance and robustness of P, PD, PI, and PID control architectures. Our theoretical findings, validated through in silico experiments using the BSim agent-based simulation platform, demonstrate the robustness and effectiveness of our multicellular PID control strategy. This innovative approach addresses critical limitations in current control methods, offering significant potential for applications in metabolic engineering, therapeutic contexts, and industrial biotechnology. Future work will focus on experimental validation in vivo and further refinement of the control models.

## 1. Introduction

Synthetic biology is an emerging field that leverages engineering principles to endow cells with new functionalities^1^. By designing innovative genetic circuits for integration into living cells, this field enables organisms to produce valuable chemicals or acquire novel capabilities, such as environmental sensing^2,3^. The applications of synthetic biology are diverse, spanning medicine, environmental science, agriculture, and the food industry. In the medical domain, for instance, researchers are developing microbial diagnostics and therapies for diseases that have eluded conventional treatments^4,5^. Furthermore, engineered microorganisms, such as yeasts, are being used to create biofuels as sustainable alternatives to fossil fuels^6^. Additionally, synthetic biology enables the production of innovative foods designed to meet specific nutritional requirements^7^.

The integration of synthetic gene networks with the native circuits of the host cell, along with the inherently nonlinear and stochastic nature of biochemical processes, can lead to unintended effects. Therefore, there is a pressing need for more reliable strategies to regulate gene expression. These strategies must ensure robust regulation of the output response while preserving the natural adaptive properties of living cells. Specifically, these strategies should maintain desired steady-state behavior even under variable conditions^8^.

The emerging field of *cybergenetics*, which merges synthetic biology with control theory, presents novel opportunities for robust regulation of biological processes. This interdisciplinary approach involves the design of synthetic feedback control architectures to ensure stable biological functioning^9,10,11^. High-performance solutions from control theory, such as nonlinear and predictive control strategies, are considered for this purpose. However, the availability of suitable biological components is limited, posing significant constraints on the feasible designs of practical controllers.

In response, the use of PID (Proportional-Integral-Derivative) controllers employing biomolecular components has gained traction due to their simplicity in implementation and their ability to provide robust regulation. These controllers leverage the antithetic motif for precise regulation and achieve a fast and damped response through derivative action. Consequently, a variety of PID controller designs have been developed, embedding PID functions directly within single cells. These designs range from those with additive dynamics of the controller actions to those incorporating nonlinear, nonseparable components^12,13,14,15,16^.

Despite the advancements, the integration of all necessary biological components within a single cell introduces challenges such as excessive metabolic burden and reduced modularity. These limitations hinder the broader applicability and scalability of the developed systems, potentially restricting their use in complex applications like immune cell engineering for cancer therapy. Thus, while the literature on embedded biomolecular PID controllers is rich with proposals, practical implementation and scalability remain critical hurdles.

Transitioning from an embedded to a multicellular approach offers a promising strategy to address the limitations associated with engineering all necessary components within one cell. In this multicellular paradigm, different functions are distributed among multiple cell populations within a microbial consortium. This division of labor can significantly minimize metabolic strain and reduce retroactivity—an unintended influence on a system caused by its interaction with another system^17,18^.

The physical separation of different biomolecular components across various cell species can alleviate the metabolic load on individual host cells. This spatial organization allows for more specialized cellular functions, which enhances the overall efficiency and effectiveness of the synthetic system^19^.

Examples of successful multicellular control designs have been documented, where engineered microbial communities coordinate their activities through the exchange of signaling molecules. These designs often utilize quorum sensing molecules to facilitate communication between different cell types, effectively closing the feedback loop necessary for robust system regulation^20,21,22^. This approach not only leverages the natural communication pathways of microorganisms but also opens new avenues for creating sophisticated, scalable, and more naturally integrated synthetic biology applications.

In this paper, we introduce a novel multicellular PID controller concept, drawing inspiration from the embedded controller design detailed in Chevalier et al. (2019)15, and building on the preliminary work on multicellular PI and PD biomolecular controllers we presented in Martinelli et al.^23,24^. Our design incorporates orthogonal quorum sensing molecules to carry out intercellular communication within a microbial consortium.

We begin by developing and detailing a mathematical model of this consortium to systematically describe its dynamics and interactions. Further, we explore the impact of varying control gains within the four types of controllers in the PID family: P, PD, PI, and PID controllers. To analyze these effects, we employ root contours, a control theoretic multi-parameter analytical technique^25^. This methodology extends the root locus analysis previously performed in Filo et al. (2023)^26^, allowing for a more comprehensive understanding of the control dynamics under different gain settings.

Our analysis provides practical guidelines for selecting the appropriate control strategy and tuning the control gains to meet specific performance criteria. To validate our theoretical findings, we conduct *in silico* experiments using BSim^27^, an agent-based realistic simulator that models the dynamics of microbial populations. These simulations help confirm the effectiveness and robustness of our proposed multicellular PID control approach under various simulated conditions.

## 2. Results

In what follows, we present the development and design of multicellular control architectures implementing a family of PID controllers. We discuss the effects of varying control gains within each controller population and validate our results through *in silico* experiments.

### 2.1. Multicellular Proportional-Integral-Derivative control architectures

We present a set of four multicellular control architectures, each comprising different engineered cell populations that implement the biological equivalents of the classical PID control schemes. Assuming the combination of all control actions, the classical PID control law can be expressed as follows:

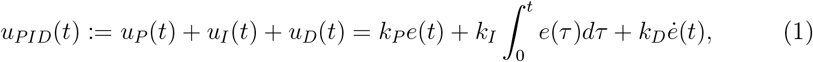

where *k*_*P*_, *k*_*I*_ and *k*_*D*_ are the so-called Proportional, Integral and Derivative control gains, respectively, and

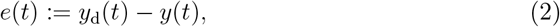

is the control error, that is, the difference between the measured output *y*(*t*) of the process to be controlled and its desired value *y*_d_(*t*). The control law in (1) is broadly and successfully employed in industrial applications due to its simplicity and ease of tunability (see for example the reference textbook28). Indeed, well-assessed techniques exist to select the values for the control gains to meet specific performance requirements.

It consists of three terms. The *Proportional* term provides an action proportional to the error *e*(*t*), causing it to decrease. However, this alone is insufficient to reduce the error exactly to zero. In theory, the error approaches zero as the proportional gain *k*_*P*_ grows larger and larger, but this is not feasible in practice. To this aim, the *Integral* action is employed to ensure robust regulation to zero, meaning that the error converges to zero regardless of constant disturbances and parametric uncertainties affecting the mathematical model. In other terms, the Integral action provides an input that compensates for constant disturbances and reference signals by integrating the error *e*(*t*). This ensures that the control signal *u*_*I*_(*t*) settles to a constant steady-state value only when the error becomes zero. Finally, by employing the time derivative of the control error *ė*(*t*), the *Derivative* action is used to make the response faster and to enhance the stability of the closed-loop system by adding damping to the response, thereby reducing excessive and undesired oscillations.

In our multicellular design, we distribute the Proportional-Integral-Derivative control actions described above among three *controller populations*. These cell populations can be combined in various ways to achieve the desired overall control signal that regulates the expression of a biological process Φ(*t*), such as a target gene, embedded in another *target population*. The modularity of our design allows for different combinations of the controller populations, enabling the realization of all possible architectures within the PID controllers’ family, namely the P, PD, PI, and PID controllers, as depicted in Figure 1a.

**Figure 1:**
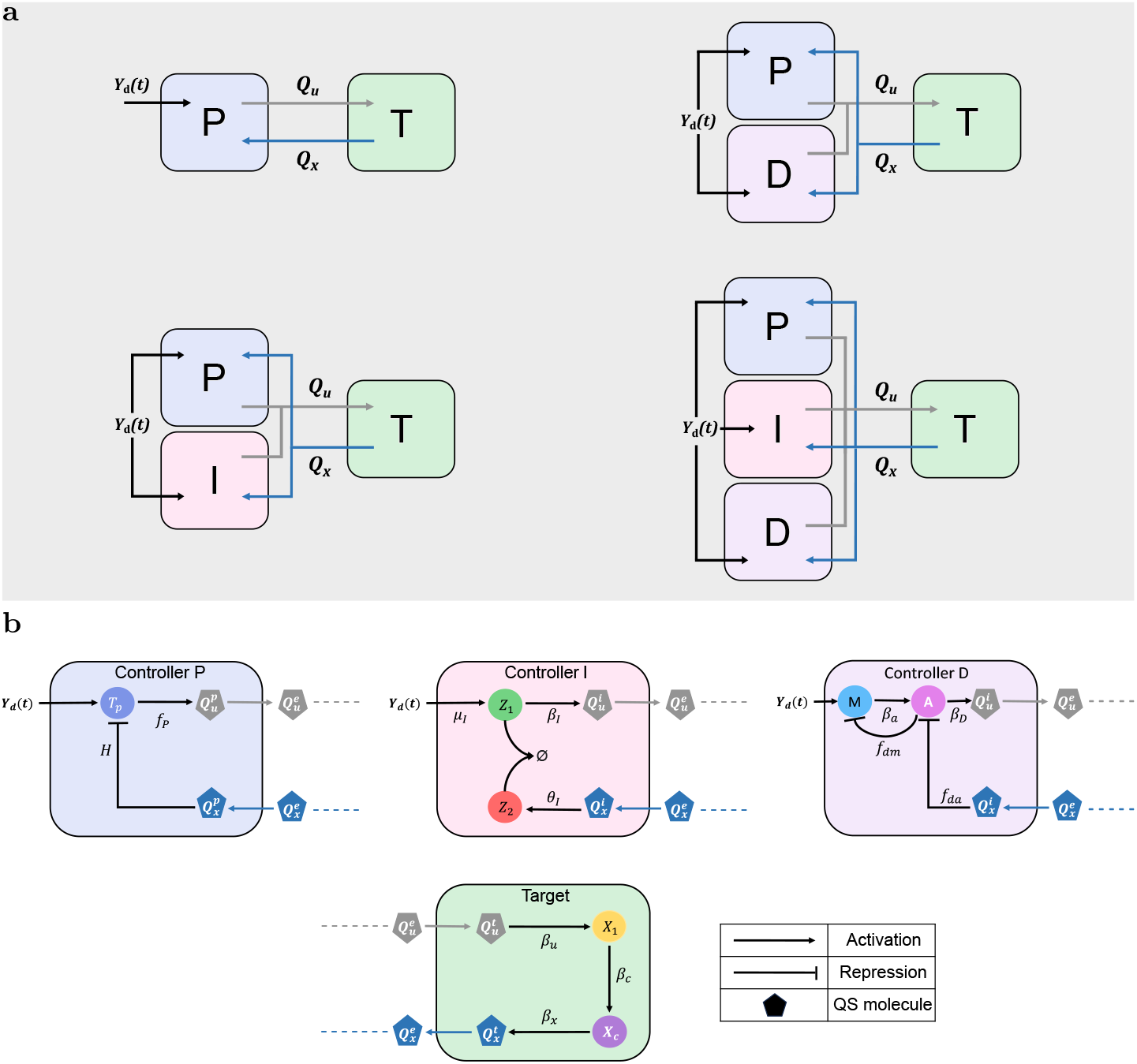
Multicellular Proportional-Derivative-Integral control architectures. The proposed multicellular control strategy consists of three controller populations that can be combined to realize one of the four multicellular PID control architectures. The molecular communication realized by the two diffusive signals *Q*_*u*_ and *Q*_*x*_ is crucial to the correct functioning of the consortium for the regulation of the biological process Φ(*t*) inside the target cells to the desired value *Y*_d_. The control input depends on the mismatch between the desired output *Y*_d_ and the measured output *Q*_*x*_ produced by the target population, that carries the information about the state of the process Φ(*t*) to the controller populations. (a) Schematic representation of the multicellular P, PD, PI and PID control architectures. Different multicellular PID family’s controllers can be obtained by combining the three controller populations. (b) Abstract biological representation of the Proportional (blue), Integral (pink) and Derivative (purple) controller cells and of the target cells (green). Each type of controller cell embeds a synthetic gene network producing the control molecule *Q*_*u*_ according to the corresponding control action. For the sake of simplicity, we assume that the process Φ(*t*) in the Target cells is a genetic network consisting of only two genes, *X*_1_ and *X*_*c*_, where *X*_1_ is directly actuated by the control signal carried by the control molecule *Q*_*u*_, and *X*_*c*_ represents the output of Φ(*t*), which in turns activates the production of the quorum sensing molecule *Q*_*x*_. Circles depict internal molecular species, while polygons represent the quorum sensing molecules. The superscripts *p, i, d* and *t* are used to designate quantities associated with the Proportional, Integral, Derivative, and Target cells, respectively, and the superscript *e* to denote quantities into the external environment. The physical meaning of all variables and parameters is reported in Table 1.

The exchange of information between the members of the consortium, enabling the cell populations to cooperate, is implemented via a pair of orthogonal quorum sensing molecules, *Q*_*u*_ and *Q*_*x*_^29^. These molecules diffuse into the environment to respectively broadcast the actuation and sensing signals, thereby closing the feedback loop. Specifically, the controllers can sense the mismatch between the reference signal *y*_d_(*t*) and the measured target output *y*(*t*), which is broadcast to them via the sensor molecule *Q*_*x*_.

**Table 1:** Table of biomolecular species’ symbols and definitions.

| Symbol | Physical meaning |
| --- | --- |
| $X_1, X_c, Z_1, Z_2, A, M$ | Cells' internal species |
| $Q_k^j, Q_k^e$ | Internal and external quorum sensing molecules |
| $\beta_u, \beta_c, \beta_x, \mu, \theta, \beta_a, \beta_m$ | Activation rates |
| $\gamma$ | Degradation rate in all cells of the consortium |
| $\gamma_a, \gamma_m$ | Active degradation rates of the derivative species |
| $\gamma_z$ | Rate constant of the annihilation reaction |
| $\gamma_e$ | External degradation rate of molecules |
| $\beta_P, \beta_I, \beta_D$ | Activation rates of the control molecule in the controller cells |
| $\eta$ | Molecules diffusion rate |
| $N$ | Number of cells in each population |
| $Y_d$ | Reference signal |

Based on this information, they produce the control molecule *Q*_*u*_ according to their designed control function.

Hence, following the principles of classical control theory, our aim is to regulate the output of the process Φ(*t*) to a desired value, ensuring that the control error *e*(*t*) converges to zero as time approaches infinity. Additionally, we require robust regulation, meaning that parametric uncertainties do not cause excessive deviation from the desired value, and that the transient response is both fast and predictable.

To assess the static and dynamic performance of the controller, we will use the following metrics: the error at steady state, *e*_∞_ = lim_*t*→∞_ |*e*(*t*)|, for static performance; and the settling time, 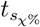, that specifies how fast the output *y*(*t*) converges inside a bound of *χ*% around the steady-state value *y*_∞_ = lim *t* → ∞*y*(*t*), and the overshoot, *o*_%_, that quantifies how large is the peak value of the oscillations in the response with respect to the steady-state value *y*_∞_, for dynamic performance.

However, due to the inherent uncertainties affecting biological systems, such as noise and parametric perturbations, asymptotic regulation, i.e., *e*_∞_ = 0, is generally not feasible. Therefore, it is more reasonable to require only bounded regulation, meaning the error norm converges below some prescribed value *ε*.

Thus, the control objective can be specified as that of achieving:

1. limt→∞ |e(t)| ≤ ε
2. 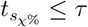 ≤ τ
3. o% ≤ ϖ

where *ε, τ* and *ϖ* are positive constants to be specified.

Despite the simplicity of the PID control law and the extensive literature and tools available for tuning its control gains in industrial applications, a systematic and straightforward approach to meet the aforementioned control specifications does not exist in the biological context we are considering here. Moreover, the multicellular architecture adds further complexity to the problem, as it requires the tuning of parameters governing the production and diffusion of signaling molecules between the cell populations. Also, while the classical PID controller family is linear, the biomolecular implementation described in this paper is inherently nonlinear making the gain tuning problem particularly cumbersome.

### 2.2. The mathematical model of the multicellular PID controllers’ family

Next, we present a mathematical model capturing the aggregate dynamics of the most comprehensive architecture, which includes all the controller populations in the consortium, namely the multicellular PID controller. This aggregate model describes the average behavior and interactions of the biochemical species within the microbial consortium, allowing us to analyze the system’s overall performance. The mathematical models for the other control architectures depicted in Figure 1a, i.e., P, PD, and PI controllers, can be directly obtained from the model of the PID controller by setting the corresponding control gains to zero.

In order to derive the aggregate dynamics, we make several simplifying assumptions. Specifically, we assume that all populations within the consortium grow and divide at the same rate. This implies that each species is also diluted at the same rate *γ*. This is a reasonable assumption when all engineered populations utilize the same microbial host. Furthermore, we assume that all populations maintain an equal number of cells, denoted by *N*, ensuring that the consortium populations are balanced in size. Building on these assumptions, it follows that the quorum sensing molecules involved in the system have uniform diffusion rates among the populations, denoted by *η*. For a comprehensive derivation of the agent-based and aggregate mathematical models, please refer to the Methods Sections 3.1 and 3.2.

In the following we use superscripts *p, i, d* and *t* to designate quantities associated with the proportional, integral, derivative, and target cells, respectively, and the superscript *e* to denote quantities into the external environment.

First, we provide the dynamics of the species produced in the target cells, which constitute the process to control, namely Φ(*t*). This process can be modeled as a network of different chemical species *X*_1_, …, *X*_*c*_, with *X*_1_ being the input species and *X*_*c*_ the output species. For simplicity, we model the process in its simplest form, as done in Briat et al. (2016)13, where the expression of *X*_*c*_ is directly influenced by *X*_1_. The abstract biological scheme of the network within the target cells is depicted in Figure 1b, where the quantity 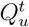 represents the actuation signal to the target cell, and the molecule 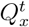 represents its measured output. Employing mass-action kinetics, the dynamics of the species produced into the targets can be written as:

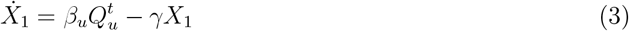

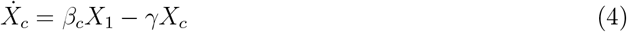

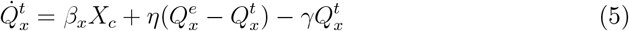

The meaning of all variables and parameters is provided in Table 1.

Next, we describe the dynamics of the control populations, whose schematic biological representation is shown in Figure 1b, each producing a contribution to the control quorum sensing molecule *Q*_*u*_. Specifically, the dynamics of the Proportional cells can be reduced to the following equation (see Section 3.3 for further details):

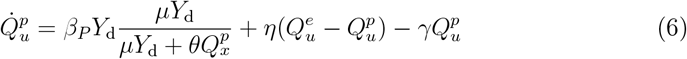

where *β*_*P*_ represents the maximal production rate of the control molecule and serves as a proportional gain.

The second contribution to *Q*_*u*_ is provided by the Integral cells, whose aggregate dynamics can be written as:

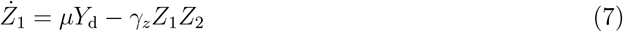

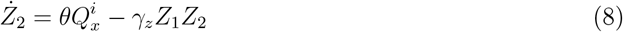

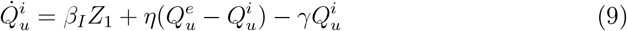

where *β*_*I*_ plays the role of an integral control gain.

The final contribution to the control molecule *Q*_*u*_ is produced by the gene network embedded within the Derivative cells, which can be modeled using Michaelis-Menten kinetics, leading to the following equations:

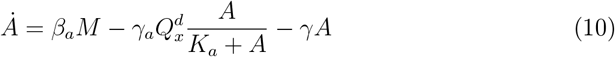

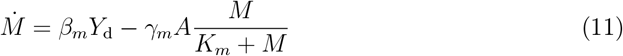

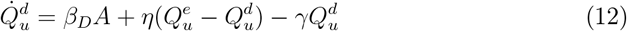

where *β*_*D*_ plays the role of a derivative control gain.

Finally, we model the dynamics of the molecules diffusing into the cells where they are not produced as:

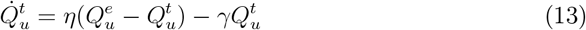

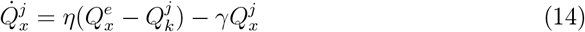

where *j* ∈ {*t, p, i, d*}, and their dynamics in the external environment as:

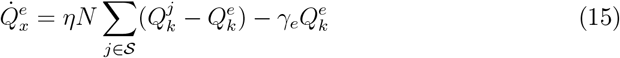

where *k* ∈ {*x, u*} and 𝒮 = {*t, p, i, d*}.

The mathematical model described by Equations (3)-(15) can be simplified by making the realistic simplifying assumptions A3-A6 on the system’s timescales (see Section 3.4). This allows us to derive a *reduced model* through timescale separation, thus obtaining:

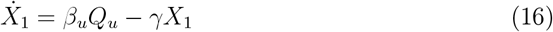

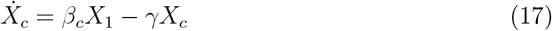

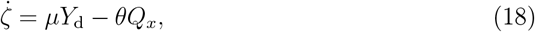

where *ζ* = *Z*_1_ − *Z*_2_. Here, the steady-state values of the quorum sensing molecules and the species *A*, respectively, are defined as:

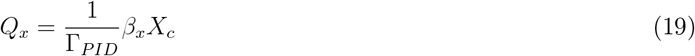

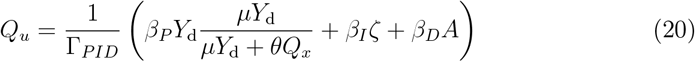

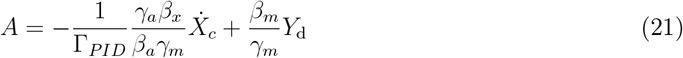

where Γ_*PID*_ = 4*γ*. Further details on the derivation of the reduced order model can be found in Section 3.4. Starting from the reduced model with PID control (16)-(18), it is possible to derive the models of the other multicellular control strategies by setting the gains of the controller populations that are not present in the consortium to zero (i.e., *β*_*I*_ = *β*_*D*_ = 0 for the P controller, *β*_*I*_ = 0 for the PD controller, and *β*_*D*_ = 0 for the PI controller). Additionally, Equation (18) is removed when the Integral cells are not present in the architecture, as is the case for the P and PD multicellular controllers.

### 2.3. Effects of Tuning Control Gains in Multicellular Control Architectures

Proportional-Integral-Derivative controllers have a different effects on the output response depending on the control actions employed and the chosen control gains. Therefore, selecting the appropriate controller is crucial to achieve the desired system performance. However, so far, differently from industrial applications, well-established tools to straightforwardly select the right values for the control gains given the desired static and transient responses are not available for the biological architecture proposed here.

Therefore, our goal is to provide guidelines for selecting the appropriate control strategy and gains to meet specific control requirements. As a result of our work, we derive analytical conditions for guiding the consortium design by evaluating the steady-state performance and the transient characteristics of the output response using multicellular P, PD, PI, and PID control architectures.

Despite the reduced model (16)-(18) simplifying the analysis of the closed-loop system dynamics, the equations remain highly nonlinear, making it difficult to evaluate how the parameters affect performance. Therefore, we performed a local analysis by linearizing the reduced model around the desired set-point, computing the closed-loop transfer function in the Laplace domain, and finally evaluating the effect of changing the control gains on the closed-loop response (see Section 3.5 for more details on the derivation). As it will be shown, this approach reveals clear relationships between control gains and closed-loop performance, which, however, hold only locally to the desired operating point and not globally. This limitation does not undermine the significance of our results, as synthetic biology applications are typically designed to operate near nominal conditions. Moreover, the validity of the control architectures we propose here will also be extensively tested in non-local conditions through in silico experiments.

The transfer function of a linear dynamical system completely describes the relationship between the input and output signals, illustrating how the closed-loop system behaves when a prescribed signal, such as the reference signal *y*_d_(*t*), is applied to it. The transfer function of the closed-loop system regulated by the complete PID controller can be expressed as follows:

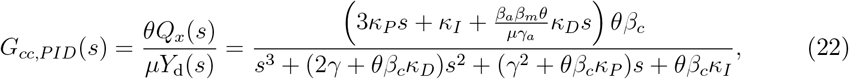

where *Q*_*x*_(*s*) and *Y*_d_(*s*) are the Laplace transform of the time-dependent functions *Q*_*x*_(*t*) and *Y*_d_(*t*), and *s* is a complex number. Here, the control gains *κ*_*P*_, *κ*_*I*_ and *κ*_*D*_ come from aggregations of the system parameters, and are defined as:

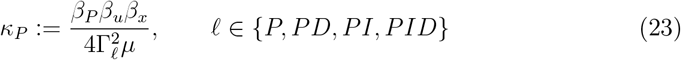

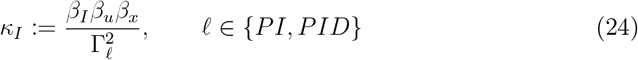

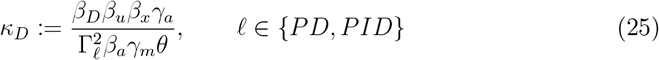

Additionally, Γ_*𝓁*_ = *Mγ* for *M* ∈ 2, 3, 4, where *M* is the number of populations comprised in the consortium. Note that the transfer functions for other control schemes, such as the P, PD, and PI control schemes shown in Figure 1a, can be derived from Equation (22) by setting the corresponding control gains to zero. For example, by setting *κ*_*I*_ = 0 and *κ*_*D*_ = 0 to obtain the P control scheme.

Next, we assessed the local performance of each control architecture leveraging the static gain of the transfer function (22) and its poles by means of the root contours analysis25 detailed in Section 3.6.

#### 2.3.1. Robust asymptotic regulation can be guaranteed only by integral action

Asymptotic regulation, meaning that the control error *e*(*t*), defined as the difference between the constant desired value *y*_d_ = *μY*_d_ and the measured output *y*(*t*) = *θQ*_*x*_(*t*), converges to zero as *t* goes to infinity, is achieved when the static gain of the closed-loop transfer function in Equation (22) is unitary.

The static gain is formally defined as the limit of the transfer function (22) as *s* tends to 0. In general, it may depends on both the control gains and the parameters of the biological system. For example, for the P controller the static gain can be computed from (22) to be 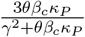. However, when the Integral controller is employed, i.e., *κ*_*I*_ ≠ 0, the static gain is always equal to 1, making it independent of the system parameters.

This means that the Integral control action, similar to the classical control law, guarantees robust asymptotic regulation. Consequently, constant perturbations from the nominal value of the parameters do not affect the output value at steady state.

Although the static gain can be equal to 1 also without the integral controller, i.e. *κ*_*I*_ = 0, as in the P and PD architectures, the choice of the control gains *κ*_*P*_ and *κ*_*D*_ that guarantee this condition depends strictly on the nominal value of the parameters of the biological system. Indeed, for the static gain to be unitary, the two control gains must satisfy the following condition:

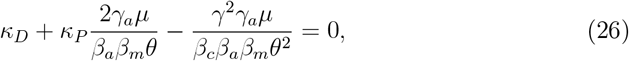

This dependence is a mathematical identity; therefore, in the P and PD cases, regulation is not robust with respect to parametric uncertainties, which are unavoidable in real biological systems. However, unlike the use of P or PD controllers, where the stability of the closed-loop system is always preserved, employing an Integral controller requires careful tuning of the control gains to ensure stability. Specifically, for the PI or PID control architectures, a unique equilibrium point exists when the control gains are adequately tuned to satisfy the condition

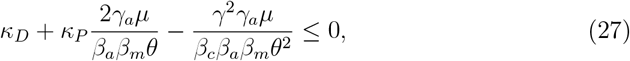

Such an equilibrium is locally exponentially stable when

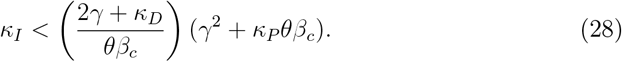

From Equation (28) it can be observed that, compared to the PI controller (where *κ*_*D*_ = 0), the PID controller extends the range of *κ*_*I*_ values for which the equilibrium remains asymptotically stable. This indicates that the Derivative action has a stabilizing effect on the closed-loop system.

The simulations reported in the right panels in Figure 2 demonstrate that when the control gains are tuned according to the aforementioned conditions, resulting in a unitary static gain, the control error at steady state of the complete model (3)-(15) remains consistent. Moreover, as will be discussed in the next section, the additional degrees of freedom in choosing the control gains for the PD, PI, and PID schemes can be further exploited to modify the transient response. This allows us to meet the control specifications for settling time 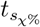 and overshoot *o*_%_..

**Figure 2:**
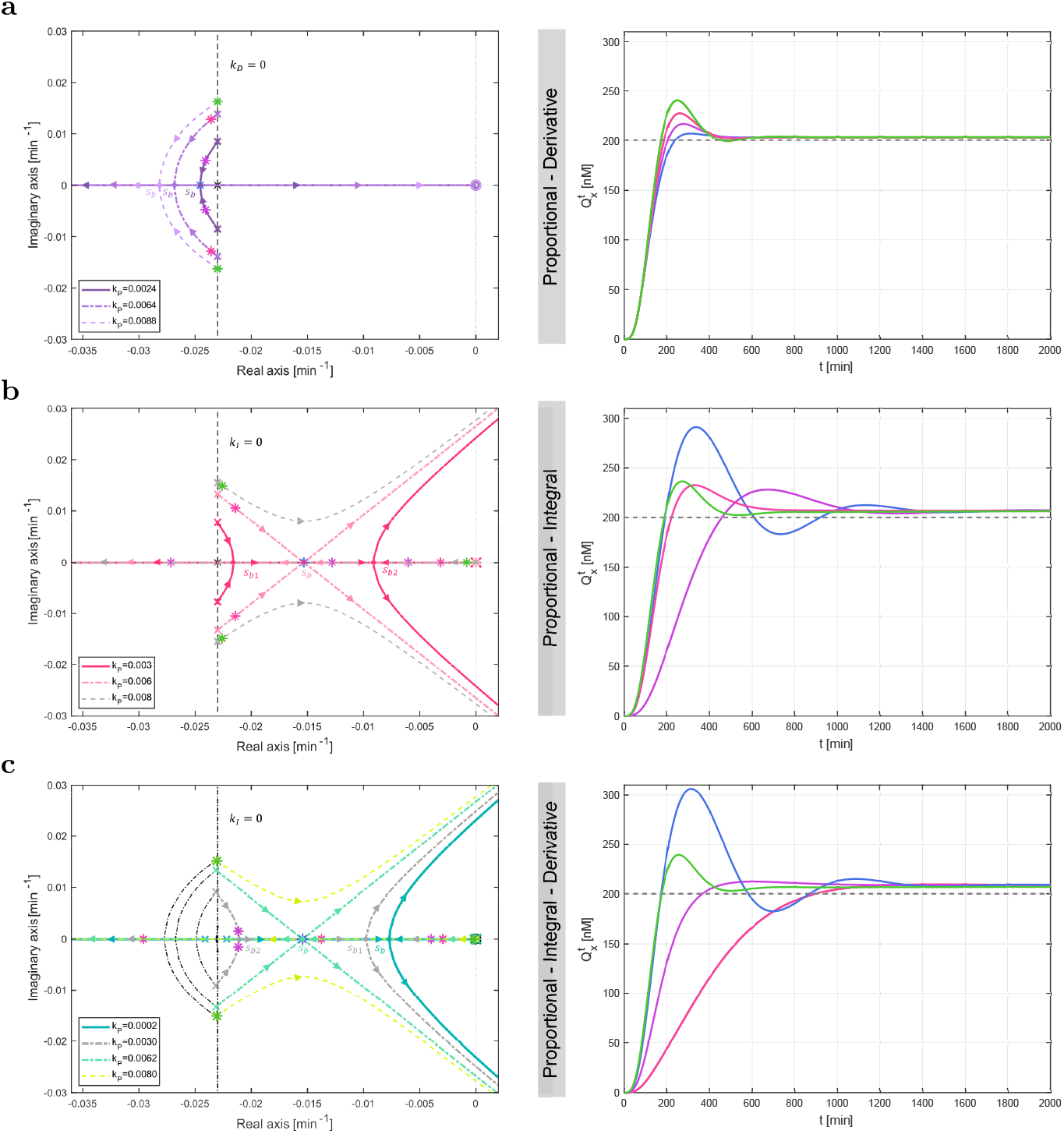
Characterization of the proposed multicellular PID family control architectures. Root contours (left panel) and time evolution of the concentration of the measured output 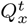 corresponding to fixed values of the control gains (right panel) when the targets are controlled by the PD, PI and PID controllers, respectively; the P controller being a special case of the PD controller when *k*_*D*_ is set to zero. The starting points of the root contours of the PD and PI architectures, respectively, lie on the root locus of *k*_*P*_ *G*_*P*_ (*s*), computed from Equation (22) when only *k*_*P*_ *≠* 0, whereas the root contours of the PID control architecture all start on the root contours of *κ*_*D*_*G*_*P D*_(*s*), computed from Equation (22) when *κ*_*P*_, *κ*_*D*_ *≠* 0. Fixing the value of *κ*_*P*_ for the PD and PI architectures, and of *κ*_*P*_ and *κ*_*D*_ for the PID architecture, allows to pick out one of the root contours. Next, changing the value of *κ*_*D*_ or *κ*_*I*_, respectively, places the closed loop poles at a specific location along the selected root contour, affecting the closed-loop transient and steady-state responses. (a) Effects of changing the derivative gain on the closed-loop poles. We selected some values of *k*_*D*_ meeting the condition (26) when only the Proportional and Derivative cells are included into the consortium. First, we considered *κ*_*D*_ = 0 (green star), that is, the Proportional controller alone. Next, we set *κ*_*D*_ *≠* 0 (pink, purple and blue star, respectively) to observe the effects of adding a derivative action on the closed-loop response. Specifically, the two real coincident poles indicated with the blue star were chosen by selecting *κ*_*D*_ meeting also condition (S75) to obtain the fastest possible response. (b) Effects of the PI controller on the closed-loop poles. We selected different values of the integral gain *κ*_*I*_ fulfilling conditions (27) and (28) (pink, purple, blue and green star, respectively). Specifically, the three coincident poles indicated with the blue star were chosen to also meet condition (S86) in order to guarantee the fastest possible output response. (c) Effects of the PID controller on the closed-loop poles. After fixing admissible values of the Proportional and Derivative gains (see condition (27)), we set some values of *κ*_*I*_ fulfilling condition (28) (pink, purple, blue and green star, respectively). Specifically, the blue star marks the three coincident real poles that are placed by choosing *κ*_*I*_ meeting also the condition derived in Section 3.6.4 to obtain the fastest possible response. The other biochemical parameters were selected as in Table S1, whereas the reference signal was set to *Y*_d_ = 60 nM.

Next, we evaluated the sensitivity of the P, PD, PI, and PID control strategies to variations in target parameters to account for parametric uncertainty in biological systems. To exploit the modularity of the proposed multicellular architecture, the values selected for each control population have been consistently used across the different schemes employing the same population. Specifically, the same values of control gain *κ*_*P*_ have been used in the simulations for the P, PD, PI, and PID control architectures; the same values of *κ*_*D*_ have been used for the PD and PID schemes; and the same values of *κ*_*I*_ have been used for the PI and PID schemes. On average, all the control architectures are able to drive the error signal near zero. However, only by including the Integral controller cells in the consortium does the steady-state error become independent of changes in *κ*_*P*_ and *κ*_*D*_ and from perturbations in the target parameters, as long as (28) is satisfied. These results are shown in Figure 3a-d, where we performed 10 simulations for each control architecture, drawing the target parameters from a normal distribution centered at their nominal value 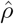 with a standard deviation of 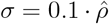. Additionally, we computed the summary statistics of the data (Figure 3e), confirming the previous analysis.

**Figure 3:**
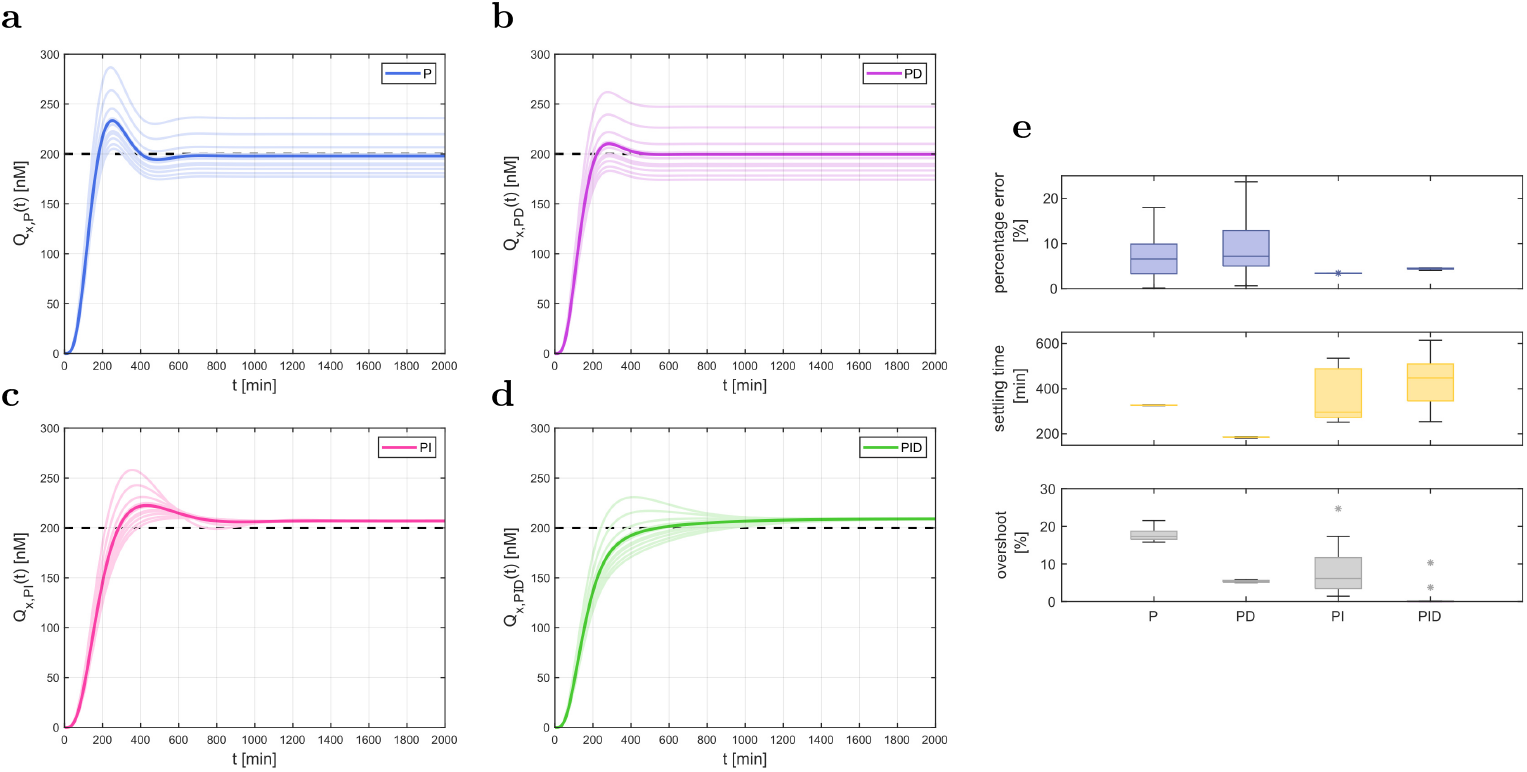
In-silico regulation experiments in Matlab. The regulation capability of the four control architectures were tested by running *in silico* experiments in Matlab. The control gains have been tuned according to Equation (27) and from the regions shown in Figure 2, fulfilling the control requirements on the transient response, and their values have been kept the same for all architecture to exploit its modularity. For each experiment the parameters of the target cells were perturbed to test the robustness of the control architecture with respect to uncertainties affecting the biological process Φ(*t*). Specifically, for each of the 10 simulations performed, the targets’ parameters were drawn from a normal distribution centered at their nominal value 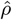 and with standard deviation 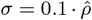. All the other parameters are fixed to their nominal values as in Table S1. All the control architectures can drive the error signal near to zero, but only when the Integral controller cells are employed the steady-state error is independent from perturbation on the parameters of the target cells. (a-d) Time evolution of the concentration of the measured output 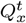 (lighter lines) when the targets are controlled by P, PD, PI and PID control actions, respectively, as the parameters of the target cells are varied. Darker lines represent the mean of these signals. (e) Box plots of the percentage steady-state error (blue), settling time at 10% (yellow) and percentage overshoot (gray) computed at each experiment of the P, PD, PI and PID control strategy, respectively. The central solid lines are the medians; the whiskers have endpoints corresponding to the minimum and maximum value excluding the outliers; the stars represent outliers. The total number of cells for each control strategy is fixed to 120, equally divided between the different cell types. Additionally, we fixed the initial concentrations of all species in the simulations to zero, assuming that each cellular population were grown separately from the others before starting the experiment.

#### 2.3.2. Derivative action makes the transient response faster

The properties of the transient response, i.e., how the output reaches its steadystate value, are closely related to the position of the closed-loop poles of the transfer function in Equation (22) in the complex plane. These poles, which are the roots of its denominator, can be either real (*s* = *a*) or complex conjugate pairs (*s* = *a* ±*jb*, where *j* is the imaginary unit). Real poles are associated with aperiodic movement, meaning a response that reaches steady-state without oscillations, while complex conjugate poles cause oscillations. The farther away the poles are from the imaginary axis, i.e., the larger the magnitude |*a*| of their real parts, the quicker their contribution to the output reaches steady-state. Therefore, the dynamical response is “dominated” by the slower poles, known as *dominant poles*, which also define the settling time of the output response, 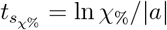. Conversely, the imaginary part of a complex conjugate pair of poles determines the period of the oscillations, and its relation with the real part determines the damping of the oscillations, and hence the maximum overshoot *o*_%_. Specifically, the damping ratio is defined as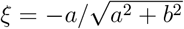 the larger the damping, the smaller the overshoot.

The position of the poles of the closed-loop transfer function (22), and in particular that of the dominant poles, can be changed by varying the control gains *κ*_*P*_, *κ*_*I*_ and *κ*_*D*_. The relationship between the control gains and the position of the poles can be studied by means of the *root contours method* (further details in Section 3.6), which allows to obtain analytical conditions on the control gains to meet the desired control specifications. Namely, the specification on the settling time 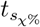 translates into requiring that the real part of the dominant poles is smaller than a prescribed value, while the specification on the overshoot *o*_%_ translates into requiring the dominant poles to be inside a triangular sector (see further details in Section 3.6). We found that the Proportional controller alone is not sufficient to make the response faster. Indeed, the real part of the dominant poles does not depend on *κ*_*P*_ as they move in the complex plane on the vertical line with coordinate 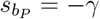 (see Figure 2a, dashed black line), where *γ* is related to the growth rate of the cells. Therefore, we need to introduce other controller populations to improve the system response.

Adding a Derivative population in the consortium to realize a multicellular PD controller, places the minimum real part achievable by the dominant pole to 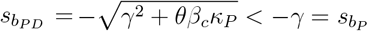, at the same time reducing the overshoot as *κ*_*D*_ increases, as also shown in Figure 2a. Thus, the PD controller can make the transient response faster and more damped than the Proportional controller alone for any value of *κ*_*P*_ *>* 0.

In contrast, when Integral controller cells are added to the consortium, that is, employing the PI and PID control architectures, the settling time worsen since all the branches of the loci are attracted toward the right-hand side. Therefore, the smallest value of the real part of the dominant poles is always greater than 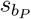 (see Figure 2c-d).

However, adding the derivative action to the PI controller improves the settling time. Indeed for any *κ*_*D*_ *>* 0 it holds that 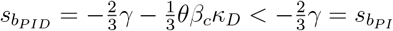.

In addition, for both the PI and PID control strategies, speeding up the response increases the percentage overshoot. This creates a trade-off between achieving a shorter settling time and minimizing the overshoot.

### 2.4. In silico experiments confirm the analysis

In this section, we present *in silico* experiments to assess the effectiveness of the proposed strategies in more realistic conditions and explore the sensitivity to parameter perturbations, and the robustness to imbalances in the composition of the consortium, of the multicellular P, PD, PI and PID control architectures.

The experiments were performed in BSim, an agent-based simulation platform designed to realistically simulate bacterial population growth dynamics [27]. By carrying out agent-based simulations in BSim, we accounted for cell geometry, cell growth and division, cell-to-cell mechanics and diffusion of molecules; additionally, we defined constraints on the host chamber.

Specifically, in all simulations we assumed the cells growing in a rescaled version of the microfluidic device employed in Shannon et al. (2020)30 and in Salzano et al. (2022)31, setting a BSim chamber of dimensions 23 *μ*m × 15 *μ*m × 1 *μ*m that can host around 120 cells, thus minimizing the computational time without losing statistical significance. For the *in silico* experimental campaign conducted here, we require the output response of the target cells to converge at steady state to the desired value with an error less than 10%, i.e., *e*_∞_ ≤ 0.1, to account for the effect of parametric disturbances affecting biological systems. Additionally, we require the transient response to settle in less than 24 hours, that is, 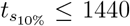, min, which is a typical timescale in controlled experimental environments [32]. Furthermore, oscillations, if present, must vanish at steady state, and the overshoot should not exceed 20% of the desired steadystate value, i.e., *o*_%_ ≤ 20%. Oscillations and excessive overshoots are undesirable in many industrial and pharmaceutical applications, such as when engineering T-cells for cancer treatment33. In all simulations, we selected the control gains according to the conditions derived in Section 2.3 and the regions depicted in Figure 2 that, for each control strategy, fulfill the prescribed control requirements.

To emphasize the modularity of the proposed control strategies, we consistently used the same values of the control gains in all control architecture, as already done in Section 2.3.1.

To evaluate the sensitivity of the system, that is, its robustness to cell-to-cell parameter variations, which naturally exist even between cells of the same species, we ran several simulations in BSim. Each time a cell split to form two daughter cells, we assigned different values to the targets’ and controllers’ parameters. Specifically, we drew the parameters from a normal distribution centered at their nominal value 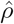 with a standard deviation *σ* = *CV* ·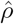, where *CV* is the coefficient of variation. The collected data showed that all the control architectures exhibit consistent performance as the intensity of the perturbation increases (Figure 4a). Specifically, the steady-state error remained almost the same as *CV* increased for all the strategies, fulfilling the desired requirement, i.e. *e*_∞_ ≤ 0.1. Regarding the settling time, the P and PD controllers showed similar performance for any of the values of the *CV* we considered, while the PI and PID controllers exhibited different behaviors as parameter perturbations increased. For *CV* = 0.15, the settling time of the PI controller increased by approximately 2.5 times compared to when there were no perturbations (*CV* = 0), while the PID controller’s settling time increased by approximately 1.2 times. Despite these variations, both strategies generally met the required condition 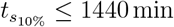, although some simulations of the PI controller exceeded this limit.

**Figure 4:**
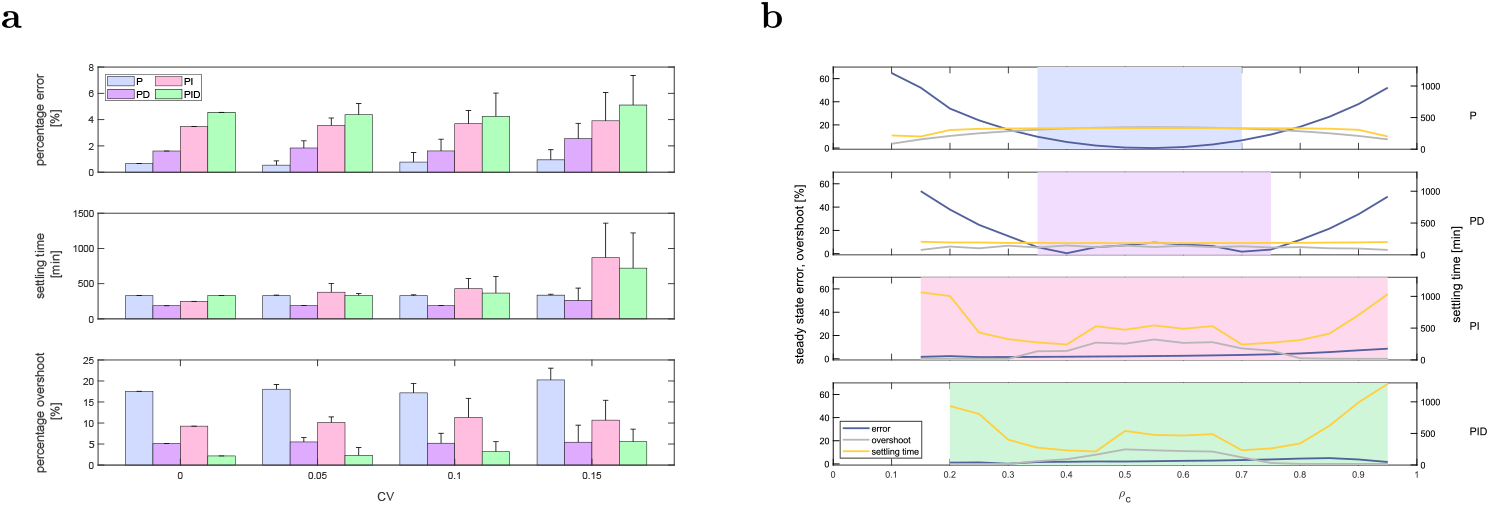
Robustness of the control architectures to parameter variations and population imbalances in BSim. All control architecture showed consistent performance as the uncertainty in the value of the biological parameters increases. Moreover, it appears that when the Integral action is employed in the consortium, the regulation capabilities of the multicellular architecture becomes more robust to imbalance between the relative number of controller and target cells. (a) Percentage error at steady state, settling time and overshoot as parameter variations increase when the targets are controlled by a P (blue bars), a PD (purple bars), a PI (pink bars) and PID (green bars) controller, respectively. For each value of the *CV* ∈ {0.05, 0.1, 0.15} we performed *n* = 20 experiments, drawing the activation parameters of the controllers and the targets from a normal distribution centered at their nominal value 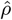 and with standard deviation 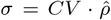. The dimensions of the BSim microfluidic chamber were different for each control strategy, in order to obtain approximately the same number of target cells at steady state for all the control architectures. The metrics were computed filtering the average targets output signal with a moving average filter of window width *W* = 60. The initial number of cells for each control strategy was equally divided between the different cell types. (b) Steady-state percentage error, settling time and overshoot of the P, PD, PI and PID control architectures as the percentage of the controller cells, *ρ*_*c*_ := *N*_*c*_*/N*, is varied, with *N* being the total number of cells in the chamber. All the simulations were performed in BSim setting the growth rate to zero in a chamber of dimensions 23 *μ*m × 15 *μ*m × 1 *μ*m hosting in total *N* = 20 cells. The shaded areas evidence the values of *ρ*_*c*_ corresponding to successful simulations, in which *e*_*∞*_ ≤ 0.1, with a settling time *t*_*s*,10%_ ≤ 1440 min and oscillations disappear at steady-state with an overshoot *o*_%_ ≤ 20%. In all experiments the reference signal is fixed to *Y*_d_ = 60 nM, while the gains are chosen as *β*_*P*_ = 0.0414 min^*−*1^, *β*_*D*_ = 0.0933 min^*−*1^ and *β*_*I*_ = 0.0002 min^*−*1^. The BSim growth and mechanical parameters were selected as in Fiore et al. (2020)^20^, while the nominal biochemical parameters were chosen as in Table S1.

Finally, the percentage overshoot *o*_%_ remained below 20% as CV increased for all control strategies, except for the P controller when *CV* = 0.15, where it slightly exceeded this upper bound. These data are reported in Figure 4a, where the relative percentage steady-state error is defined as:

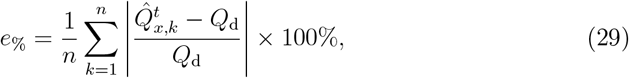

with 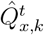 being the value of 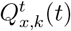, averaged over the last 600 min, of the *k*-th experiment, 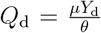 the desired value of *Q*_*x*_, and *n* the total number of experiments for each value of *CV*.

Next, we considered the presence of imbalances in the consortium composition, which might arise from the different metabolic loads associated with the expression of each control action, affecting the growth rate of the cells.

To test robustness to imbalances, we performed several simulations in BSim, each with a different relative number of cells across the populations while neglecting cell growth and division. The presence of the Integral cells in the consortium provides robustness (shaded areas in Figure 4b), and ensures also a smaller error at steadystate. However, the settling time 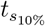 worsens for the PI and PID controllers, although it remains under 1440, min. Meanwhile, the percentage overshoot *o*_%_ stayed almost constant and below 20% for all the control strategies.

For each value of *ρ*_*c*_ the steady-state error has been evaluated as follows:

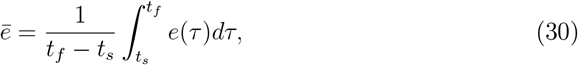

where, 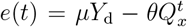 is the error signal, *t*_*s*_ = 1440 min, and *t*_*f*_ is the simulation time.

## 3. Discussion

In this paper, we introduced a novel multicellular Proportional-Integral-Derivative (PID) control strategy for regulating biological processes. Our comprehensive mathematical analysis and model derivation together with in silico experiments demonstrate the robustness and efficacy of this approach, highlighting several key findings and potential applications, as well as areas for future improvement.

Our work successfully demonstrated a modular design approach, where PID control actions are distributed among three distinct controller populations within a consortium. This modularity allows for flexible combinations to achieve desired control architectures (P, PD, PI, and PID) tailored to specific biological processes. The mathematical models developed for these control architectures, derived from the aggregate dynamics of the system, enable systematic analysis of their performance. This flexibility is crucial for tailoring the control strategies to various biological contexts, ensuring precise and effective regulation.

We found that, despite the nonlinear nature of the biomolecular implementation we presented, the inclusion of Integral controller cells in the consortium guarantees robust asymptotic regulation as is the case in classical PID controllers. This robustness is evident as the steady-state error remains independent of parameter variations, provided specific conditions are met. Such robust regulation is essential for maintaining system stability despite the inherent parametric uncertainties in biological systems. This result is particularly relevant for applications where precise control is paramount, such as the development of reliable devices and networks in synthetic biology and metabolic engineering.

Furthermore, our analysis showed that adding Derivative action to the control strategy significantly enhances the transient response by reducing the settling time and dampening oscillations. This improvement in transient performance is critical for applications requiring quick and stable responses. However, it is also noteworthy that while the PI and PID control architectures improve steady-state accuracy, they introduce trade-offs in transient performance, particularly in terms of increased settling time. These trade-offs must be carefully managed to balance the overall system performance.

The in silico experiments conducted using the BSim platform validated our analytical results in the presence of more realistic effects including cell growth, division, and molecular diffusion.

Our multicellular PID control strategy has broad potential in applications. In synthetic biology, it can be applied to various scenarios where precise control of gene expression is essential, such as in metabolic engineering for optimizing the production of pharmaceuticals or biofuels. In biomedical engineering, our approach can be extended to therapeutic contexts, such as engineering T-cells for cancer treatment, where maintaining a stable and precise response to environmental cues is crucial for effective therapy. Additionally, in industrial biotechnology, where microbial consortia are employed for chemical production or bioremediation, our control strategy can enhance process stability and yield by mitigating the effects of environmental and parametric uncertainties.

Despite the promising results, there are areas for future improvement. Experimental validation in vivo is necessary to confirm the practical applicability and effectiveness of our control strategies in real biological systems. To this aim we are exploring the use of multi-chamber bioreactors that could be used to grow different cell populations in different chambers and combine them in a mixing chamber where densities could be carefully controlled using strategies such the ratiometric control techniques presented in34,31. Further additional work could focus on developing more sophisticated models that capture additional layers of biological complexity, such as stochastic effects, spatial heterogeneity, and dynamic interactions among cell populations. Implementing advanced optimization algorithms for tuning control gains could further improve the performance and robustness of the control architectures, making them more adaptive to varying biological conditions. Exploring the scalability of our control strategies for larger and more complex consortia, as well as their integration with other synthetic biology tools and frameworks, will be crucial for broader application and adoption.

## Methods

Detailed methods are provided in the online version of this paper and include the following:

- RESOURCE AVAILABILITY
- Lead Contact and Materials Availability
- Data and Code Availability
- METHOD DETAILS
- Derivation of the agent-based mathematical model
- Aggregate dynamics of the consortium,
- Proportional cells’ network dynamics
- Derivation of the reduced order models of the PID controller family
- Root contours analysis

## Acknowledgements

M.d.B. wishes to acknowledge support from the Scuola Superiore Meridionale under the funding provided to the Area on Modeling and Engineering Risk and Complexity. D.F. wishes to acknowledge support by the European Union Next Generation EU, under PRIN 2022 PNRR, Project “Control of smart microbial communities for wastewater treatment”.

## Author Contributions

M.d.B. developed the initial concept and framework of the study, with D.F. contributing to refining the research questions and methodology. V.M. with support from D.F. and D.S. derived and refined the mathematical model. V.M. and D.S. coded and conducted the in-silico experiments in Matlab and BSim. V.M., D.F. and D.S. with support from M.d.B. analyzed the data. The original draft of the manuscript was prepared by V.M., with significant input and revisions from D.F. and the other authors. All authors have read and agreed the final version of the manuscript.

## Declaration of generative AI and AI-assisted technologies in the writing process

During the preparation of this work the author(s) used ChatGPT4o in order to revise the grammar, eliminate repetitions and typos. After using this tool, the authors reviewed and edited the content as needed and take full responsibility for the content of the publication.

## Supplemental information

Document S1. Table S1, Figures S1–S2 and Video S1.

Table S1. Simulation parameter values used in simulations, related to Figures 2-4.

Figure S1. Abstract biological implementation of the PID controller.

Figure S2. Root locus of the multicellular P controller.

## METHODS

### Lead Contact and Materials Availability

Further information and requests for resources should be directed to and will be fulfilled by the Lead contact, Mario di Bernardo.

### Data and Code Availability

The code used for all simulations is available at https://github.com/Vittoria96/MulticellularPIDcontrol.

## METHOD DETAILS

### 3.1. Derivation of the agent-based mathematical model

We derive the mathematical model of the most comprehensive multicellular ProportionalIntegral-Derivative control architecture, specifically the multicellular PID controller, describing the agent-based dynamics35 of the consortia. A schematic biological representation of the proposed control architecture is depicted in Figure S1. The primary function of the controller populations is to regulate the concentration of a biological process Φ(*t*) inside the target cells. The process Φ(*t*) consists of a network of various chemical species *X*_1_, …, *X*_*c*_ interconnected through a series of reactions, in which *X*_1_ is the input species and *X*_*c*_ the output species.

Following the methodology described in Briat et al. (2016)13 and in Chevalier et al. (2019)15, we model the process Φ(*t*) in its simplest form, whereby *X*_1_ directly promotes the expression of *X*_*c*_. The synthesis of the species *X*_1_ is driven by the control quorum sensing molecule *Q*_*u*_. Conversely, the sensor quorum sensing molecule *Q*_*x*_ is synthesized proportionally to the concentration of the output species *X*_*c*_. To describe the dynamics of *X*_1_ and *X*_*c*_ within each target cell, we employ mass-action kinetics, leading to the following equations:

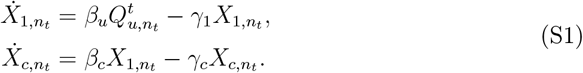

where *n*_*t*_ = 1, …, *N*_*t*_, and *N*_*t*_ is the number of target cells in the consortium. Additionally, in Equation (S1), *β*_*u*_ and *β*_*c*_ represent the production rates of *X*_1_ and *X*_*c*_, respectively, and *γ*_1_ and *γ*_*c*_ are their degradation rates. The target cells also produce the sensing molecule *Q*_*x*_, whose dynamics is given by:

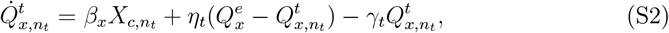

where *β*_*x*_ represents the activation rate of 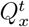, while *γ*_*t*_ denotes its degradation rate, which is assumed to be uniform across all target cells. Furthermore, 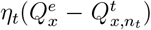 describes the diffusion dynamics of the sensor molecule across the target cells’ membrane. This process is typically considered to be linear36, with *η*_*t*_ being the diffusion rate.

The control action is carried out via the control molecule *Q*_*u*_, produced by the three bacterial populations, namely the Proportional, Integral and Derivative controller cell populations. The Integral controller cells contain a synthetic network based on the antithetic motif presented in Briat et al. (2016)13. This circuit consists of two molecular species, denoted as *Z*_1_ and *Z*_2_, that bind together forming an inert complex. Specifically, *Z*_1_ promotes the production of the molecule 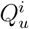, and in turn *Z*_1_ is produced proportionally to the reference signal *Y*_d_. Conversely, the activation of *Z*_2_ is proportional to the sensor molecule *Q*_*x*_. Note that, the binding reaction between *Z*_1_ and *Z*_2_ effectively acts as a comparator between the output of the target process and the reference signal37. The overall dynamics of the antithetic network inside Integral cells can be described as:

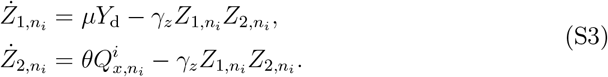

with *n*_*i*_ = 1, …, *N*_*i*_, *N*_*i*_ being the number of cells in the Integral population. Additionally, 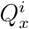 is the concentration of the molecule *Q*_*x*_ into the Integral cells, *μ* and *θ* denote the activation rates of *Z*_1_ and *Z*_2_, respectively, and *γ*_*z*_ is the annihilation rate.

Note that we assumed the cell growth dynamics to be much slower than the other chemical reactions of interest in the cells, implying that the controller species *Z*_1_ and *Z*_2_ only degrade forming the inert complex, as similarly assumed in Briat et al. (2016)13. A more realistic model can be derived by assuming nonzero degradation rates of *Z*_1_ and *Z*_2_^38^).

The integral controller population contribute to the overall control action by producing the control molecule 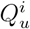 proportionally to *Z*_1_. Specifically, the dynamics of the control molecule *Q*_*u*_ into the Integral cells can be written as:

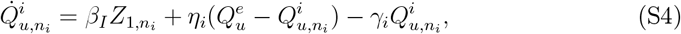

where *β*_*I*_ is the activation rate of 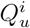 and plays the role of an integral gain, *γ*_*i*_ is the degradation rate of the molecules in the Integral cells and *η*_*i*_ is the diffusion rate from the external to the internal integral cells’ environment.

The second contribution to the production of *Q*_*u*_ comes from the Proportional controller cells. The Proportional controllers rely on a gene regulatory network similar to the one presented in Chevalier et al. (2019)15, based on two transcription factors competing for the same promoter of a gene named *T*_*p*_. Assuming that the protein expressed by *T*_*p*_ has a fast decay rate, we can neglect its dynamics (more details can be found in Section 3.3) reducing the dynamics of the Proportional controllers network to that of the molecule *Q*_*u*_, which can be described by:

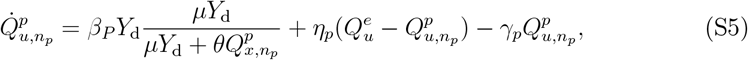

where *n*_*p*_ = 1, …, *N*_*p*_ and *N*_*p*_ is the number of Proportional controllers in the consortium. The parameters *μ* and *θ* are incorporated into the activation function of 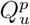 to maintain a consistent notation with the antithetic network as in Chevalier et al. (2019)15. Furthermore, the activation rate of 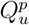, namely *β*_*P*_, is a tunable parameter which plays the role of a proportional gain, *η*_*p*_ is the diffusion rate across the proportional controllers’ membrane, and *γ*_*p*_ is the degradation rate of the quorum sensing molecules into all the Proportional cells. Note that, the activation rate of 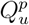, which contains the reference signal *Y*_d_, can be demonstrated to be a nonlinear function of the error using similar arguments to those reported in15 for embedded PID control.

The final contribution to the control action is exerted by the Derivative cell population. These cells embed a gene network designed to carry out a derivative action on the Target cells. The network we considered here is inspired by the motif present in Ma et al. (2009)39 and in Chevalier et al. (2019)15. Specifically, we designed a derivative network constituted by two biomolecular species *A* and *M*, whose dynamics can be written as:

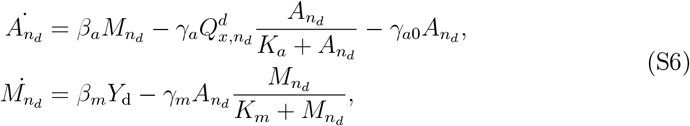

where *n*_*d*_ = 1, …, *N*_*d*_, with *N*_*d*_ denoting the number of Derivative cells. Additionally, 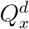 is the concentration of the sensor molecule *Q*_*x*_ in the Derivative cells, *β*_*a*_ and *β*_*m*_ represent the activation rates of *A* and *M*, respectively, and *γ*_*a*0_ is the degradation rate of *A*. The active degradation of *A* and *M* occurs through enzymatic reactions modeled via Michaelis-Menten functions40 with constants *K*_*a*_ and *K*_*m*_, and their rates are denoted with *γ*_*a*_ and *γ*_*m*_, respectively. Assuming *K*_*a*_ ≪ 1 and *K*_*m*_ ≪ 1, that is, the enzyme are saturated, it is possible to demonstrate that the network described by Equation (S6) implements an approximate time derivative of the output *X*_*c*_^15^. The Derivative cells contributes to the production of the control molecule *Q*_*u*_, whose dynamics inside these cells is given by:

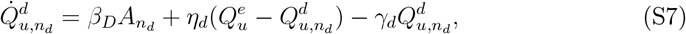

where *β*_*D*_ denotes the activation rate of *Q*_*u*_, and plays the role of a derivative gain, *η*_*d*_ is the diffusion coefficient across the derivative cells’ membrane and *γ*_*d*_ is the degradation rate of the quorum sensing molecules.

Next, we model the dynamics of the quorum sensing molecules within cells not involved in their production as:

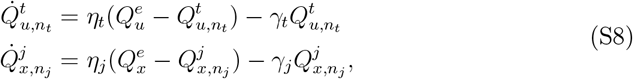

where *n*_*j*_ = 1, …, *N*_*j*_ for *j* ∈ {*t, p, i, d*} is the number of cells in the *j*-th population, 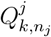 and 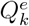 are the concentrations of *Q*_*x*_ and *Q*_*u*_ into the target, proportional, integral and derivative cells and into the external environment, respectively, and *γ*_*j*_ represents their degradation rate.

Finally, we complete the agent-based model by providing the description of the dynamics of the quorum sensing molecules into the external environment:

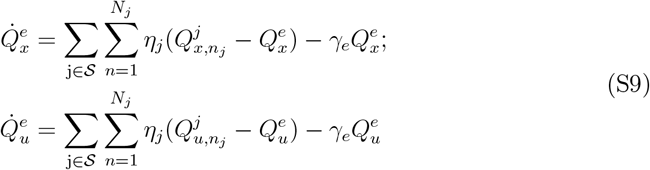

where 𝒮 = {*t, p, i, d*}, and *γ*_*e*_ is the degradation rate of the quorum sensing molecule into the external environment.

### 3.2. Aggregate dynamics of the consortium

Here, we derive the aggregate dynamics of the multicellular PID architecture presented in Section 2.2, describing the evolution of the average concentration of all the chemical species within the microbial consortium. The aggregate model formulation is based on the agent-based model described by Equations (S1)-(S9).

Assuming the average concentrations of the non-diffusive species (i.e. *X*_1_, *X*_*c*_, *Z*_1_, *Z*_2_, *A* and *M*) follow the same dynamics, their aggregate dynamics straightforwardly follow from the agent-based model. Regarding the quorum sensing molecules, their agent-based dynamics are given as follows:

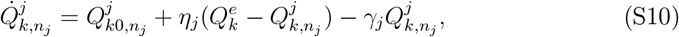

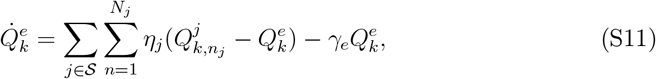

where 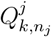 and 𝒮 = {*t, p, i, d*} are the concentrations of the signaling molecules inside the *n*_*j*_-th cell of the respective population, 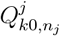 are the activation rates, and 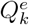 are the concentrations of *Q*_*x*_ and *Q*_*u*_ into the environment. To derive the aggregate dynamics of the diffusing molecules, we make the following simplifying assumptions:

*A1. All cells in the consortium have the same growing and dividing rates*.

*A2. The populations are balanced, that is, every population is composed by the same number of cells, i.e. N*_*t*_ = *N*_*p*_ = *N*_*i*_ = *N*_*d*_ = *N*.

Assumptions A1 and A2 imply that the quorum sensing molecules have the same diffusion rate *η* across the cell membrane. Thus, by defining 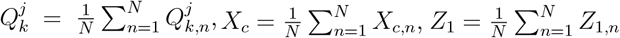 and 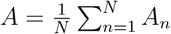, we can derive the evolution of the average concentration of the signaling molecules inside each cell population as:

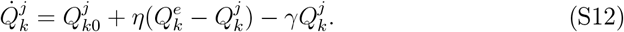

In addition, we can recast Equation (S11) as:

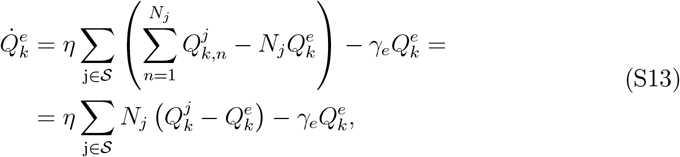

Using Assumption A2, Equation (S13) becomes:

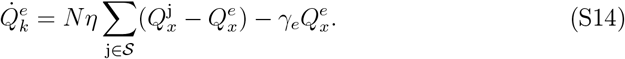

### 3.3. Proportional cells’ network dynamics

In our model, the proportional cells’ dynamics were derived based on an assumption made by Chevalier et al. (2019)15, which allows the derivation of the proportional action from an experimental realization they proposed. In the following, we describe the same procedure applied to the proportional cells of the multicellular architecture to derive the dynamics in Equation (S5). The gene network comprises two transcription factors *T*_*x*_ and *T*_*y*_, whose activation is proportional to the concentrations of the sensor molecule *Q*_*x*_ and of the reference signal *Y*_d_, respectively, competing for the promoter of a gene *T*_*P*_, whose dynamics can be written as:

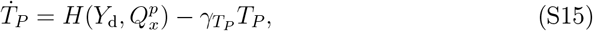

where 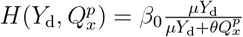 derives from a scenario in which transcription can occur only when *T*_*y*_ (activated by *Y*_d_) is bound to the promoter of *T*_*P*_, while it is repressed when *T*_*x*_ (activated by *Q*_*x*_) is bound. Additionally, the quorum sensing molecules dynamics is given by:

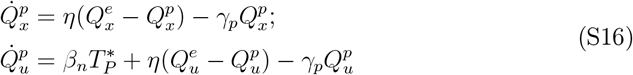

where 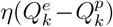 for *k* ∈ {*x, u*} describes the diffusion of the molecules through the cell membrane, and 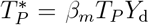 is the active form of the transcription factor produced from the gene *T*_*P*_ when *T*_*y*_ binds to its promoter. Moreover, *β*_*n*_ is the production rate of 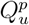, while *γ*_*p*_ is the degradation rate of the quorum sensing molecules into the Proportional cells. By applying a time-scale separation on Equation (S15), choosing 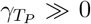, and performing the same manipulations as in Chevalier et al. (2019) ^15^, the overall dynamics of the network in the Proportional cells can be described by only Equation (S16), thus recasting the dynamics of *Q*_*u*_ as:

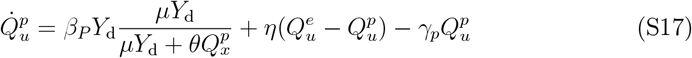

where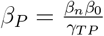.

### 3.4. Derivation of the reduced order models of the PID controller family

We derive the reduced version of the multicellular PID model by making the following realistic simplifying assumptions:

*A3. The quorum sensing molecules diffuse much faster than they degrade, that is η* ≫Γ_*𝓁*_, *with* Γ_*𝓁*_ = *Mγ for 𝓁* ∈ {*P, PI, PD, PID*} *and M the number of populations in the consortium. This assumption was showed to hold in different models parameterized from in vivo experiments*^*20,36*^.

*A4. The annihilation process between the species Z*_1_ *and Z*_2_ *of the antithetic motif in* (S3) *is fast enough, that is, the value of γ*_*z*_ *is sufficiently high in order for* 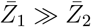 *to be at steady-state*^*41*^.

*A5. The enzymatic reactions in* (S6) *take place near saturation, which is a common assumption*^*40*^ *and implies K*_*a*_ ≪ 1 *and K*_*m*_ ≪ 1.

*A6. The dynamics of the derivative network in* (S6) *are sufficiently faster than those of the target process* (S1), *allowing for a time-scale separation to be performed between the dynamics of the process to control and the derivative dynamics*^*15*^.

Based on these simplifying assumptions, in the following we derive the reduced order model of the multicellular PID control architecture, from which the P, PI and PD reduced models can be easily obtained.

#### 3.4.1. Quasi-steady state analysis of the dynamics of the quorum sensing molecules

Starting from Assumption A3, we make a quasi-steady state assumption on the dynamics of the quorum sensing molecules, that is, by imposing 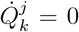 and 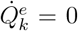 in Equations (S12) and (S14), respectively, with *k* ∈ {*u, x*} and *j* ∈ 𝒮, where 𝒮 = {*t, p, i, d*}. Thus obtaining:

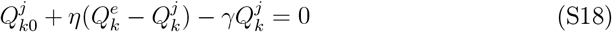

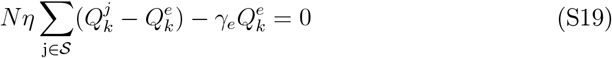

where *N* is the number of cells in each population, assumed to be balanced. From (S18), the expression of 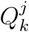 can be written as:

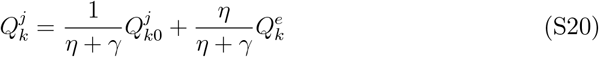

Next, substituting (S20) in (S19), we obtain:

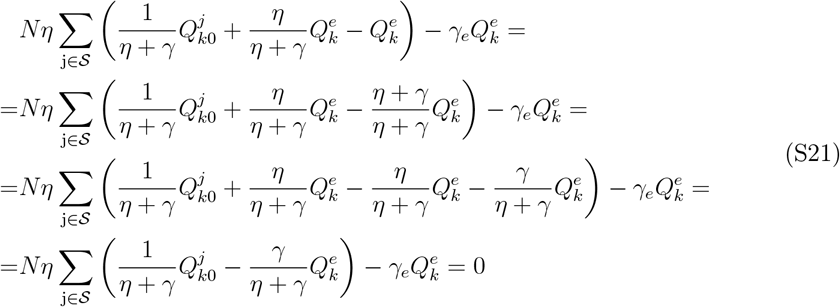

From the assumption that *η* ≫ Γ_*𝓁*_, it also follows that *η* ≫ *γ*, thus we can write:

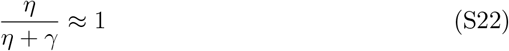

Hence, substituting the approximation (S22) in (S21), we obtain:

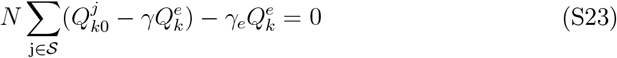

and, separating the sum and using the definition of Γ_*𝓁*_ := *Mγ* where *M* is the number of populations in the consortium, the previous equation becomes:

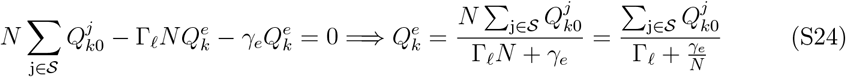

Assuming that the number of cells *N* is very large, we finally obtain:

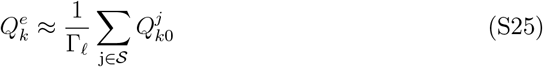

Next, we substitute (S25) in (S20), thus obtaining:

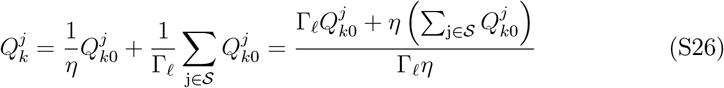

Given that:

- 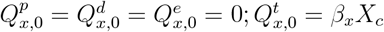
- 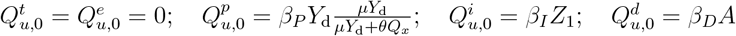

and substituting the expression rate of the quorum sensing molecules *Q*_*x*_ and *Q*_*u*_ in each population in (S26) using again the Assumption A3, at the end we have at steady-state:

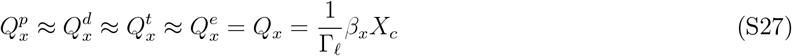

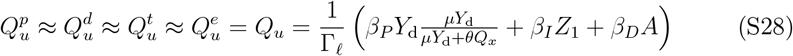

The steady-state values of the quorum sensing molecule for each of the presented topologies can be derived from (S27) and (S28) by setting to zero the dilution rates and the control gains of the populations that are no part of the selected control architecture.

#### 3.4.2. Singular perturbation analysis of the integral dynamics

Based on Assumption A4, we perform a singular perturbation analysis on Equation (S3), choosing as perturbed parameter the annihilation rate *γ*_*z*_. Firstly, we introduce the variable *ζ* = *Z*_1_ − *Z*_2_ in place of *Z*_2_, transforming equations (S3) into:

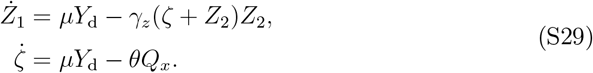

where *Q*_*x*_ is given by (S27). Next, we apply the strong binding assumption (Assumption 3). In case that the targets are regulated by only Integral controllers, 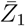 and 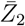 can be computed to be:

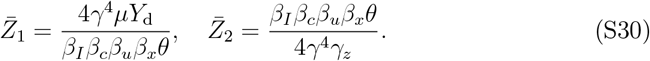

Thus, the strong binding assumption can be written as:

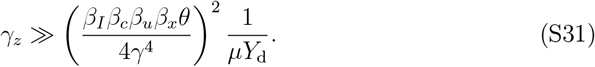

In the limit of strong binding, *Ż*_1_ can be considered at steady-state42, and dilating time *τ* := *γ*_*z*_*t* the dynamics of the variable *Z*_1_ can be described by:

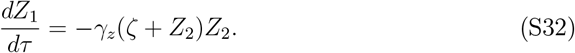

Equation (S32) admits two solutions, i.e. *Z*_2_ = −*ζ* and *Z*_2_ = 0. Computing the Jacobian of (S32) it is easy to verify that only *Z*_2_ = 0 is a stable equilibrium, and thus an admissible solution, so the system (S29) becomes:

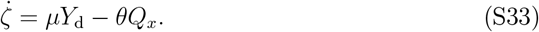

#### 3.4.3. Time-scale separation between derivative and target dynamics

Based on Assumptions A5 and A6, we perform a time-scale separation between the dynamics of the derivative network and the one of the controlled process. Applying Assumption A5, that implies *K*_*a*_ ≪ *A* and *K*_*m*_ ≪ *M*, and the quasi-steady state analysis in Section 3.4.1, Equation (S6) becomes:

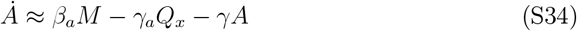

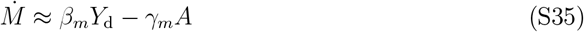

Taking the time-derivative of Equation (S34) and substituting *Q*_*x*_ with its expression (S27), solving it for 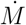 and substituting the resulting expression into (S35), we obtain:

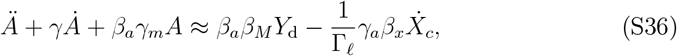

with *𝓁* ∈ {*PD, PID*} according to the multicellular controller of interest. At this point, we apply a Laplace transform on (S36), obtaining:

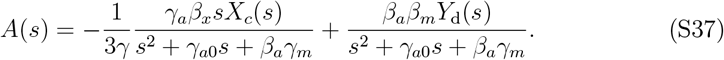

As similarly done in Chevalier et al. (2019)15, we next assume *β*_*a*_*γ*_*m*_ ≫ |*s*^2^ + *γ*_*a*0_*s*|, that is equivalent to 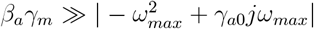 with *ω*_*max*_ the upper bound of the frequency content of *X*_*c*_(*t*), which means there is a time-scale separation between the dynamics of *A*(*t*) and *X*_*c*_(*t*). In such a way, the denominator of the *X*_*c*_(*s*), which is a band-pass transfer function, becomes influential only at high frequencies and *A*(*s*) is approximately proportional to *sX*_*c*_(*s*). Under this assumption, Equation (S37) becomes:

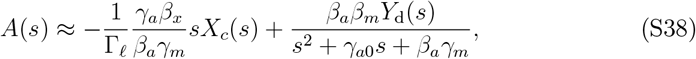

which can be rewritten in the time domain as:

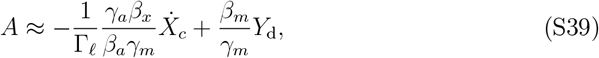

where the inverse Laplace transform of the transfer function between *Y*_d_(*s*) and *A*(*s*) has been computed considering *Y*_d_ as a succession of step functions and neglecting the brief transient in the time domain signal of *A* due to the step changes in *Y*_d_^15^. It is now easy to see that, under the assumption of time-scale separation, *A* is an approximated form of the derivative of the error defined as *e*(*t*) := *μY*_d_ − *θQ*_*x*_, due to the direct relationship between *X*_*c*_ and *Q*_*x*_.

#### 3.4.4. Reduced order model

Based on Assumptions A1-A6, the multicellular PID model presented in Section 2.2 can be reduced to:

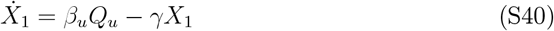

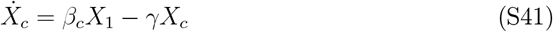

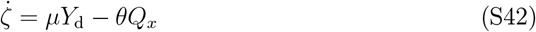

where *Q*_*x*_, *Q*_*u*_ and *A* are given in (S27), (S28) and (S39), respectively, with Γ_*𝓁*_ for *𝓁* ∈ {*P, PI, PD, PID*} depending on the multicellular control topology of interest. Additionally, Equation (S42) is removed when the the Integral cells are not present in the architecture, i.e. for P and PD multicellular controllers.

### 3.5. Derivation of the transfer functions through linearization of the dynamics around the set-point

To compute the transfer function of the reduced order system, we start from the model (S40)-(S42) linearized about the steady-state 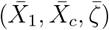 corresponding to the desired set-point value 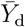, obtaining:

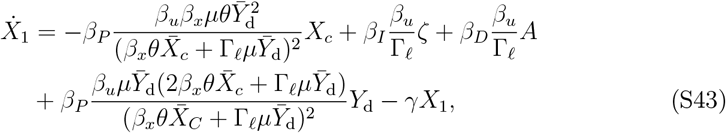

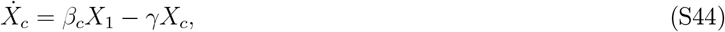

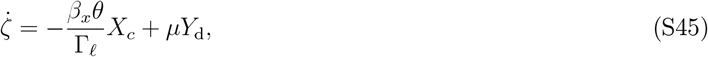

where *A* is given by (S39), *𝓁* ∈ {*P, PI, PD, PID*} and *β*_*P*_, *β*_*I*_, *β*_*D*_ are nonzero only if the Proportional, Integral and Derivative cells are comprised in the consortium, respectively. Additionally, Equation (S45) is removed for P and PD multicellular controllers. Next, we recast the terms in Equation (S43) as:

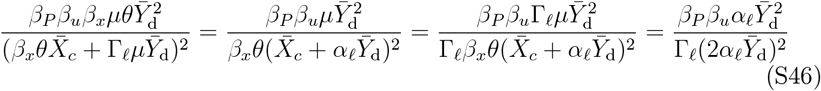

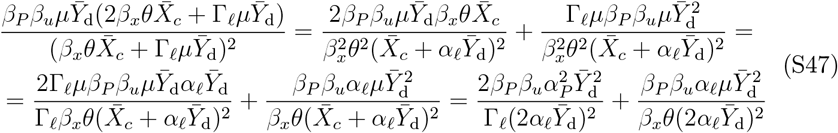

where 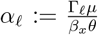 and we substituted 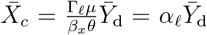, which is the desired steadystate value of *X*_*c*_(*t*). We then define:

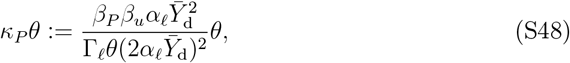

thus obtaining:

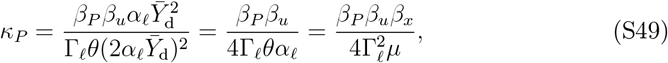

where we substituted the expression of 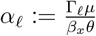. The first term of equation (S47) can be then rewritten as:

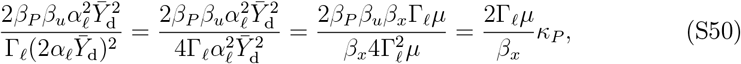

while the second term of equation (S47) becomes:

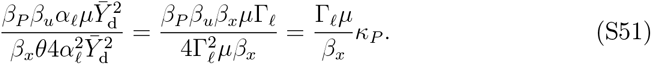

Summing (S50) and (S51) we obtain:

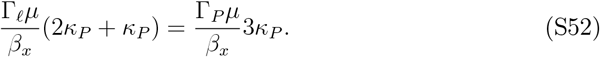

Defining 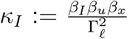 and 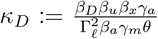 and performing some manipulations on the quantities in 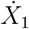, the Laplace transform of (S43) can be therefore computed to be:

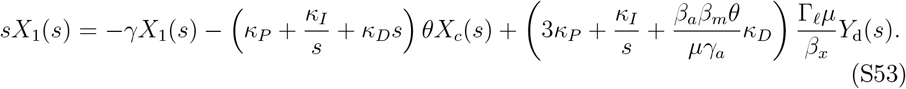

where we substituted 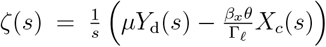 and 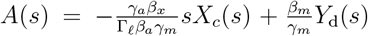, that is, the Laplace transform of (S39). Additionally, the Laplace transform of 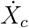 (S44) is given by:

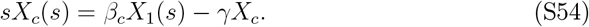

If *f* (*s*) is the transfer function between *X*_1_ and *X*_*c*_, that is:

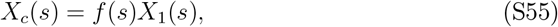

then we have:

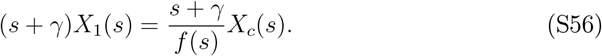

Defining 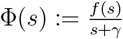, from (S53) we finally obtain:

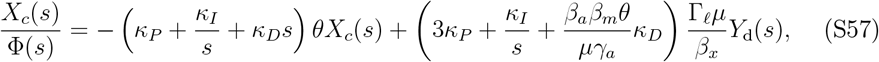

which can be recast as:

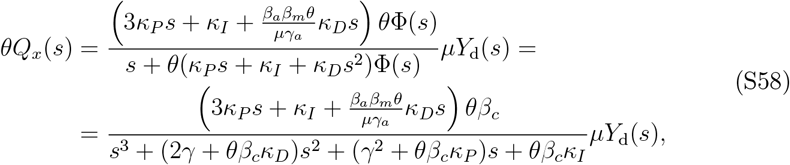

where we substituted 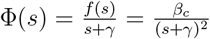 and 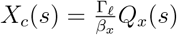. Starting from (S58), the transfer functions of the P, PI, PD and PID multicellular control architectures can be derived by setting to zero the control gains of the populations that are not included in the consortium and for Γ_*𝓁*_ with *𝓁* ∈ {*P, PI, PD, PID*}, where the control gains are defined as:

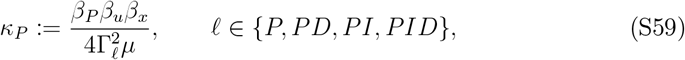

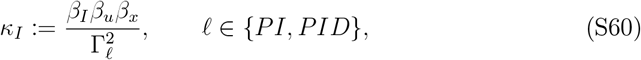

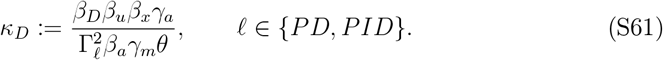

Note that since the P and PD controller guarantee zero steady-state error only when condition (26) is met, the linearization is valid only for those values of the control gains and with unperturbed dynamics.

### 3.6. Root contours analysis

To support the tuning of the gains, we apply the root locus and root contour analysis described by Golnaraghi et al. (2017)25 to the P, PD, PI and PID control architectures. In so doing, we derive analytical conditions on the control parameters parameters of the controllers’ multicellular implementations to adjust their static and dynamic performance.

#### 3.6.1. P controller: root locus

Here, we describe the procedure to compute the root locus of the Proportional control architecture. Starting from the transfer function (S58), by setting *κ*_*D*_ = *κ*_*I*_ = 0, we obtain the transfer function:

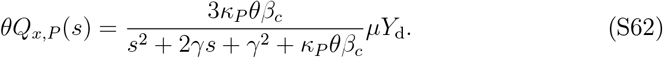

Next, Equation (S62) can be recast as:

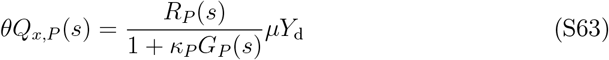

where:

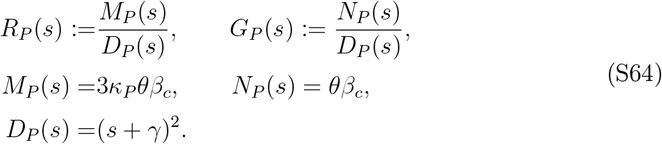

In this form, the numerator *R*_*P*_ (*s*) in (S63) does not affect the position of closed-loop poles in the complex plane. Therefore, we can study how the proportional gain *κ*_*p*_ affects the dynamics of the closed-loop system by just studying the root locus of the function:

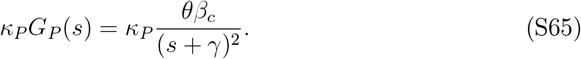

Specifically, *G*_*P*_ (*s*) has two real poles:

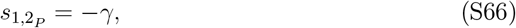

thus the root locus has two branches starting at −*γ* and going to infinity along the vertical axis defined by (S66). Therefore, the effect of *κ*_*P*_ is to increase the imaginary part of the closed-loop poles, and thus increasing the oscillations in the response of the system. Instead, the real part, and thus the settling time, remains unchanged. Moreover, the system is always stable varying *κ*_*P*_ (see Figure S2).

#### 3.6.2. PD controller: root contours

To analyze the proposed multicellular PD controller, we first derive the closed-loop transfer function from Equation (S58). For *κ*_*I*_ = 0, it can be computed to be:

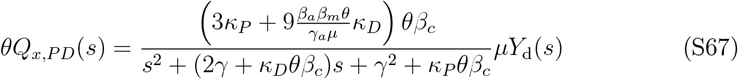

As done for the multicellular Proportional controller in Section 3.6.1, we recast the transfer function of the PD Controller (S67) as follows:

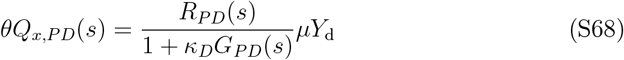

where:

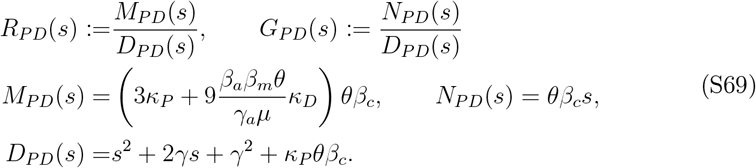

Since in this case we have to study the effects of varying two parameters on the closedloop poles, namely *κ*_*P*_ and *κ*_*D*_, we assess the local performance of the PD architecture using the root contours method25, that is a multi-parameter method to construct the root locus of the closed-loop function. The first step consists into setting the denominator of the closed-loop transfer function (S67) to zero:

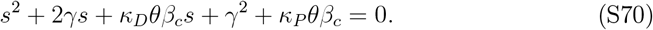

Next, we set *κ*_*D*_ equal to zero, and dividing both sides of the equation (S70) by *s*^2^ + 2*γs* + *γ*^2^, we obtain:

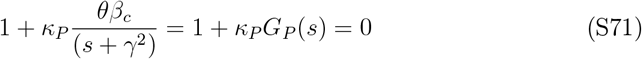

which is the same denominator as in the case of the Proportional Controller. Thus, the root locus when *κ*_*D*_ = 0 is the same as in Figure S2. At this point, we restore the value of *κ*_*D*_ and recast the equation (S70) to obtain:

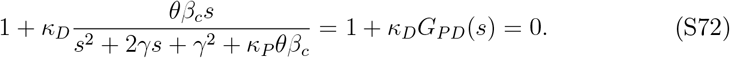

By fixing *κ*_*P*_, it is possible to move the poles of the input-output transfer function in Equation (S72) on the line *s* = −*γ*. Thus, each fixed value of *κ*_*P*_ allows to pick out one of the root contours whose starting points lie on this line. For instance, the different curves in the left panel of Figure 2a correspond to different chosen values of the proportional gain. After selecting one of the root contours, changing the value of *κ*_*D*_ places the closed loop poles at a specific location along it, altering the closed-loop response.

All the root contours have a similar qualitative behaviour. Specifically, their branches all belong to left-half plane, making the closed-loop system stable for any value of the control gains. Additionally, they start from two complex conjugates poles on the root locus of *κ*_*P*_ *G*_*P*_ (*s*) (S65), which setting *κ*_*D*_ = 0 can be computed to be:

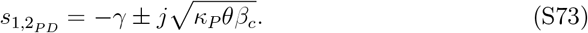

Then, the real and imaginary parts of the closed loop poles decrease increasing *κ*_*D*_ until ℑ{*s*_1_} = ℑ{*s*_2_} = 0 and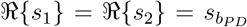, that is the most negative real part achievable from the dominant pole that allows to achieve the smallest settling time. Specifically, the breaking point can be computed to be:

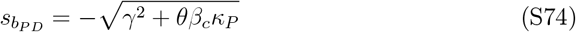

The value of *κ*_*D*_ needed to place the poles at 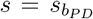 on each root contour can be then derived using the Magnitude Condition25, and it is given by:

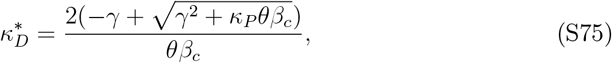

Hence, the position of 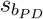 can be modified by choosing different values of *κ*_*P*_. Thus, the PD controller makes the response arbitrarily fast.

#### 3.6.3. PI controller: root contours

Here, we apply the root contours method25 to the PI control system, in order to analyze how the biomolecular control gains influence the closed-loop poles of its transfer function. First, we compute the closed-loop transfer function by setting *κ*_*D*_ = 0 in Equation (S58), thus obtaining:

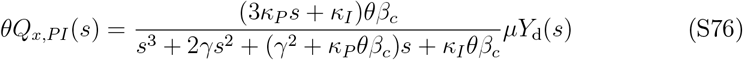

Next, we recast the transfer function by recasting Equation (S76) as:

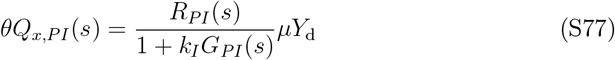

where:

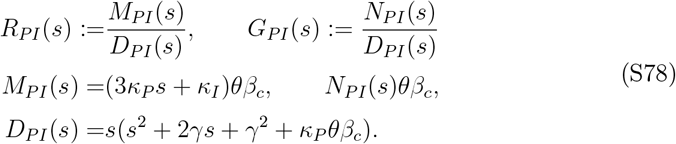

To apply the root contours method, we set the denominator of the transfer function (S76) to zero:

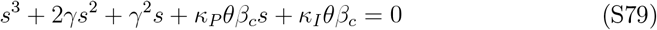

Next, we consider *κ*_*I*_ = 0, obtaining again the same input-output transfer function *κ*_*P*_ *G*_*P*_ (*s*) as the Proportional Control architecture (S65). Restoring then *κ*_*I*_, we can rewrite equation (S79) as:

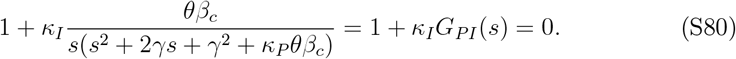

The starting points of the root contours of *G*_*PI*_(*s*) will thus lie on the root locus of *G*_*P*_ (*s*). Specifically, each root contour has three branches starting from the open-loop poles of *G*_*PI*_(*s*), which can be computed to be:

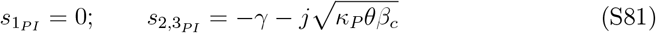

Then, each branch will extend to infinity.

Differently from the P and PD architectures, the equilibrium point is positive only if:

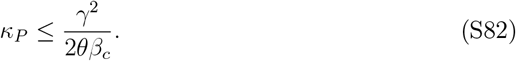

Hence, we have an upper bound on the value of *κ*_*P*_, limiting the region where the starting points of the PI root contours can lie.

Additionally, the closed-loop system is locally asymptotically stable only if:

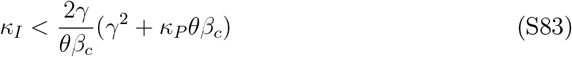

Therefore, the higher *κ*_*P*_ is, the higher the integral gain that causes loss of stability. The range of *κ*_*I*_ values guaranteeing stability at steady state can be widened either by choosing fast-dividing cells or by reducing the strength of the promoters induced by *Q*_*x*_ (smaller value of *θ*).

By fixing the value of *κ*_*P*_ fulfilling the condition in Equation (S82), it is possible to change the position of the poles of the input-output transfer function along the line *x* = −*γ*, thus selecting one of the root contours (e.g., curves in the left panel of Figure 2b correspond to different fixed values of *κ*_*P*_). After fixing *κ*_*P*_, the closed-loop poles can be placed along the selected root contour by changing the value of *κ*_*I*_. Specifically, when *κ*_*I*_ = 0, all the root contours start from a dominant real pole at zero and two complex conjugate poles with negative real parts. As the value of *κ*_*I*_ increases, the dominant pole becomes more negative, while the real part of the complex conjugate poles increases as the imaginary part decreases.

Specifically, by fixing a root contour with 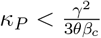 (see Figure 2b for *κ*_*P*_ = 0.003), the complex poles starting from *κ*_*P*_ *G*_*P*_ (*s*) collide at the breaking point *s*_*b*1_ and become real, while the real pole value decreases as *κ*_*I*_ increases. The system then has three real poles until two of them collide at the breakaway point *s*_*b*2_, becoming complex conjugates and dominating the response. The mathematical expression of the breakpoints can be derived as follows:

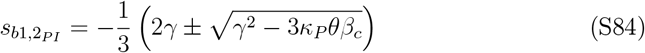

Note that, under the above condition on *κ*_*P*_, the dominant pole of each root contour cannot be placed further to the left than the breakaway point 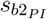, for the following value of the integral gain:

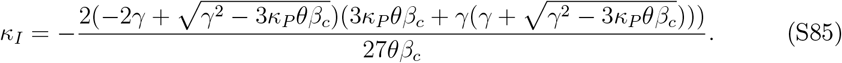

In order to minimize the value of 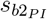, and thereby the settling time, we fix the proportional gain exactly to 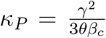 (see Figure 2b for *κ*_*P*_ = 0.006). This causes the two breakpoints *s*_*b*1_ and *s*_*b*2_ to overlap at 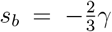, making the system have one real pole and two complex poles for any value of *κ*_*I*_. Specifically, the closed-loop poles collide and exchange at *s*_*b*_ for the following value of *κ*_*I*_ computed through the Magnitude Condition:

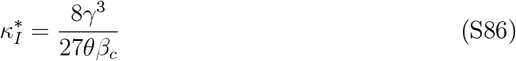

that is, the value to choose to obtain the fastest settling time. Finally, for 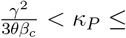 (see Figure 2b for *κ*_*P*_ = 0.008), the system has a real pole and two complex poles that do not collide at any break-point but exchange dominance on the response at *s*_*b*_.

#### 3.6.4. PID controller: root contours

As for the other proposed control architectures, here we apply the root contours method25 to investigate the effects of varying three control gains of the PID multicellular implementation on the closed-loop response. Specifically, the transfer function of the multicellular PID control system is given by Equation (S58), and it can be recast as:

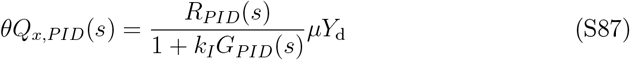

where:

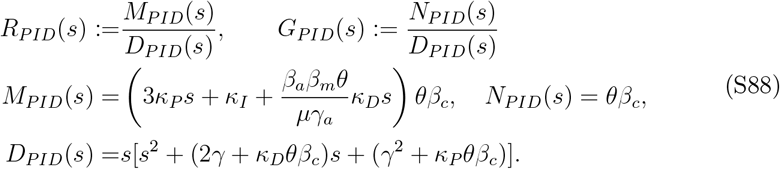

Next, we set:

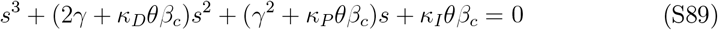

in order to derive the root contours by first setting the value of all but one of the control gains to zero, and then restoring one of them at each step. Specifically, for *κ*_*D*_ = *κ*_*I*_ = 0 we obtain:

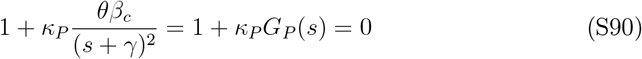

that contains the same input-output transfer function as the Proportional Controller (S65). Next, restoring *κ*_*D*_, Equation (S89) can be recast as:

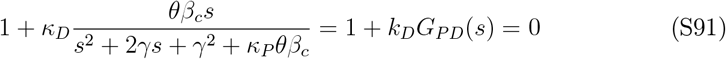

which embodies another already studied input-output transfer function, that is *G*_*PD*_(*s*) (S72). Thus, the starting points of the root contours restoring *κ*_*D*_ will lie on the root locus of *G*_*P*_ (*s*). Finally, with the values of all the gains different from zero, Equation (S89) can be computed to be:

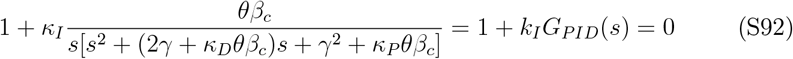

Therefore, restoring *κ*_*I*_ last, the root contours of *G*_*PID*_(*s*) will start on one of the root contours of *G*_*PD*_(*s*) derived in Section 3.6.2, selected by fixing a value of *κ*_*D*_, and whose start points are on the root locus of *G*_*P*_ (*s*). In particular, the starting points of the root contours are the poles of *G*_*PID*_(*s*), which can be computed to be:

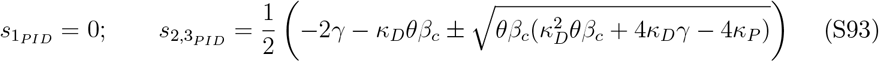

Thus, each root contour has three branches starting at the poles and extending to infinity.

Additionally, we found that the equilibrium point of the closed loop system is positive, and thus admissible, only if:

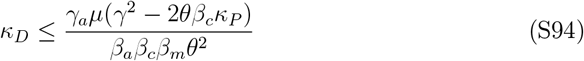

which identifies the prohibited regions for each PID contours’ starting points to lie. The root contours show again that in the presence of the Integral cells in the consortium the system remains stable only until:

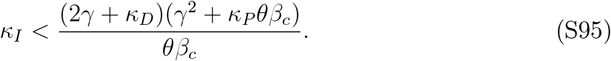

This stability range can be widened either by selecting a higher value of *κ*_*P*_ or a higher value of *κ*_*D*_, along with using fast-dividing cells or reducing the strength of the promoter induced by *Q*_*x*_ in the integral cells.

Given the above conditions, 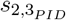 in (S93) are real only if 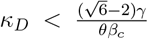 and 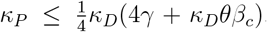, or 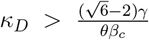 and 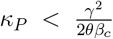 (e.g. see Figure 2c for *κ*_*P*_ = 0.0002); then, increasing the value of *κ*_*I*_, the closed loop-system has three real poles until the breakaway point:

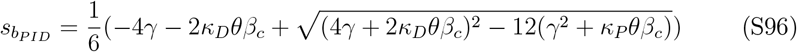

which corresponds to the minimum value achievable by the dominant poles on each root contour, where two real branches become complex. In the limit condition 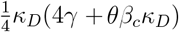, the breakaway point becomes 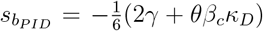, which is its leftmost reachable value when the closed-loop poles are real, obtainable for 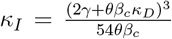 even if the infimum value in this case can be computed to be 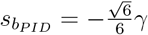. Next, the starting points of the root contours, that is, the open-loop poles 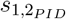, are complex conjugates if 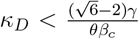 and 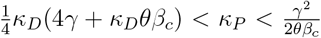, as shown in Figure 2c for *κ*_*P*_ = 0.0030. Under these conditions, the complex branches become real at the breaking point 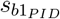 and the real branches become complex at the breakaway point 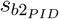, respectively, whose expressions are given by:

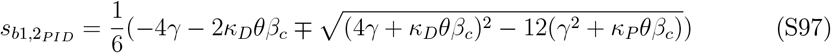

until 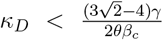 and 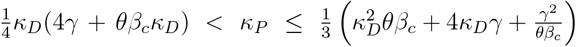, or 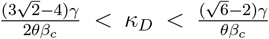 and 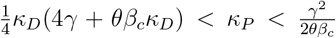. When *κ*_*P*_ is chosen exactly as 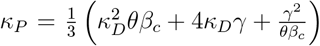 as in Figure 2c for *κ*_P_ = 0.0062, the closed loop system has again a real pole and two complex poles exchanging their dominance in *s*_*b*_, which derives by the overlapping of *s*_*b*1_ and *s*_*b*2_. In this case, the breakpoint cannot be placed further to the left than 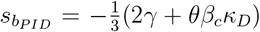 in correspondence of 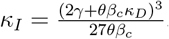, although its infimum value can be computed to be 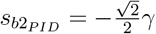 Finally, for 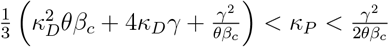 (Figure 2c for *κ*_P_ = 0.0080), the root contour has no breakpoints and the closed loop system has always one real pole and two complex conjugate poles, whose real part is the same in correspondence of *s* = *s*_*b*_, that is, the value of *s* in correspondence of which the real part of the complex poles becomes closer to zero with respect to the real pole.

## Supplemental Information

### List of Tables

**S1 Simulation parameter values used in simulations**

### List of Figures

**S1 Abstract biological scheme of the proposed PID control architecture**

**S2 Root locus of the proposed multicellular P control architecture**

**Table S1:** Nominal biochemical parameters for the designed multicellular feedback Proportional-Integral-Derivative control architectures.

| Parameter | Value | Reference |
| --- | --- | --- |
| $\beta_u$ | $0.06 \text{ min}^{-1}$ | [20] |
| $\beta_c$ | $0.1 \text{ min}^{-1}$ | [15] |
| $\beta_x$ | $0.03 \text{ min}^{-1}$ | [20] |
| $\mu$ | $1 \text{ min}^{-1}$ | [15] |
| $\theta$ | $0.3 \text{ min}^{-1}$ | [15] |
| $\gamma_z$ | $0.01 \text{ nM}^{-1} \text{ min}^{-1}$ | [15] |
| $\beta_a$ | $1.5 \text{ min}^{-1}$ | [15] |
| $\gamma_a$ | $1.5 \text{ min}^{-1}$ | [15] |
| $K_a$ | 1 nM | [15] |
| $\beta_m$ | $0.4167 \text{ min}^{-1}$ | [15] |
| $\gamma_m$ | $1.5 \text{ min}^{-1}$ | [15] |
| $K_m$ | 1 nM | [15] |
| $\gamma$ | $0.023 \text{ min}^{-1}$ | [20] |
| $\gamma^e$ | $0.0023 \text{ min}^{-1}$ | // |
| $\eta$ | $2 \text{ min}^{-1}$ | [20] |

**Figure S1:**
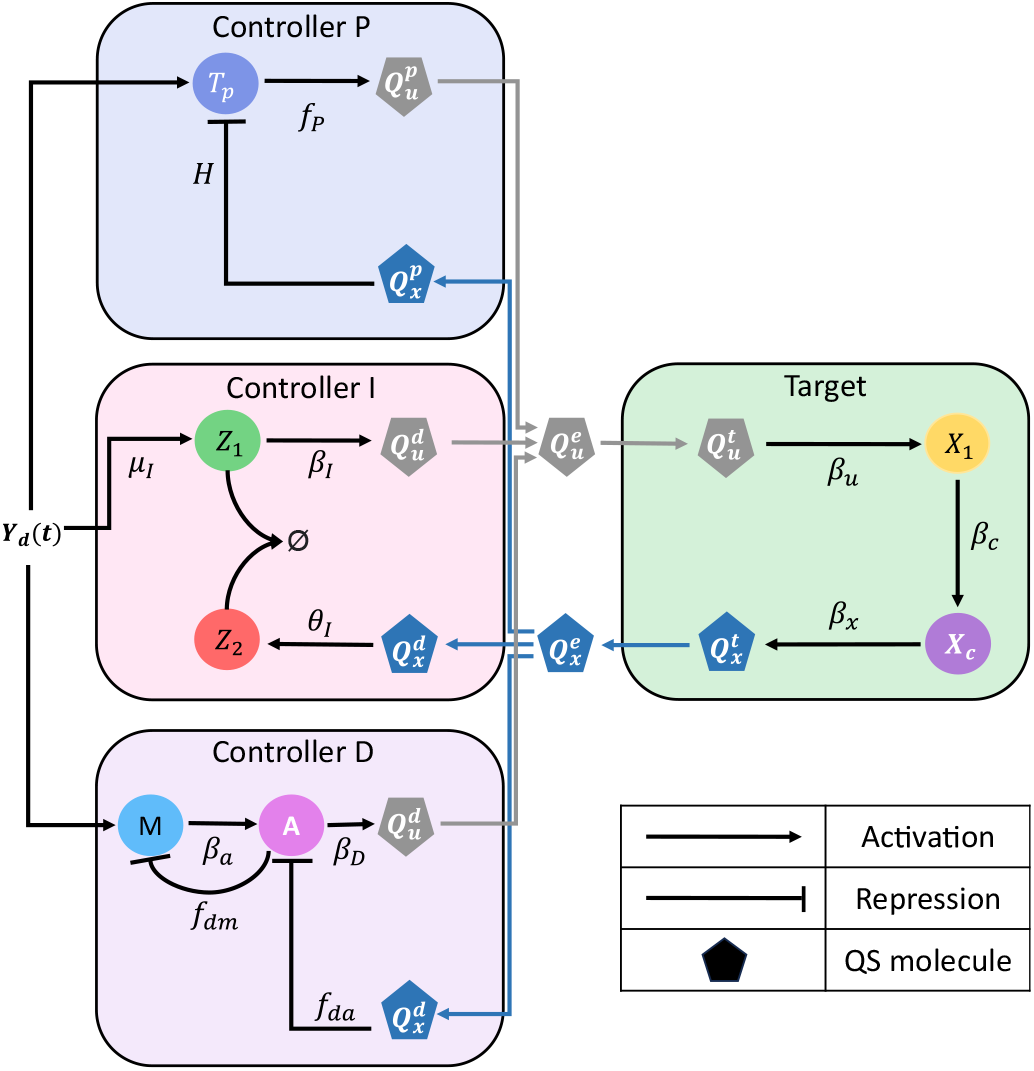
Abstract biological scheme of the proposed PID control architecture. Abstract implementation of the designed multicellular PID controller. The output of the target process embedded in the target cells is the quorum sensing molecule *Q*_*x*_, produced proportionally to the output *X*_*c*_. The input of this process is the gene *X*_1_, which is directly actuated by the control quorum sensing molecule *Q*_*u*_. Each controller population evaluates the control error by comparing the reference signal *Y*_d_(*t*) and the output signal carried by *Q*_*x*_, thus contributing to the overall production of *Q*_*u*_. Circles represent internal molecular species, while polygons represent the signaling molecules

**Figure S2:**
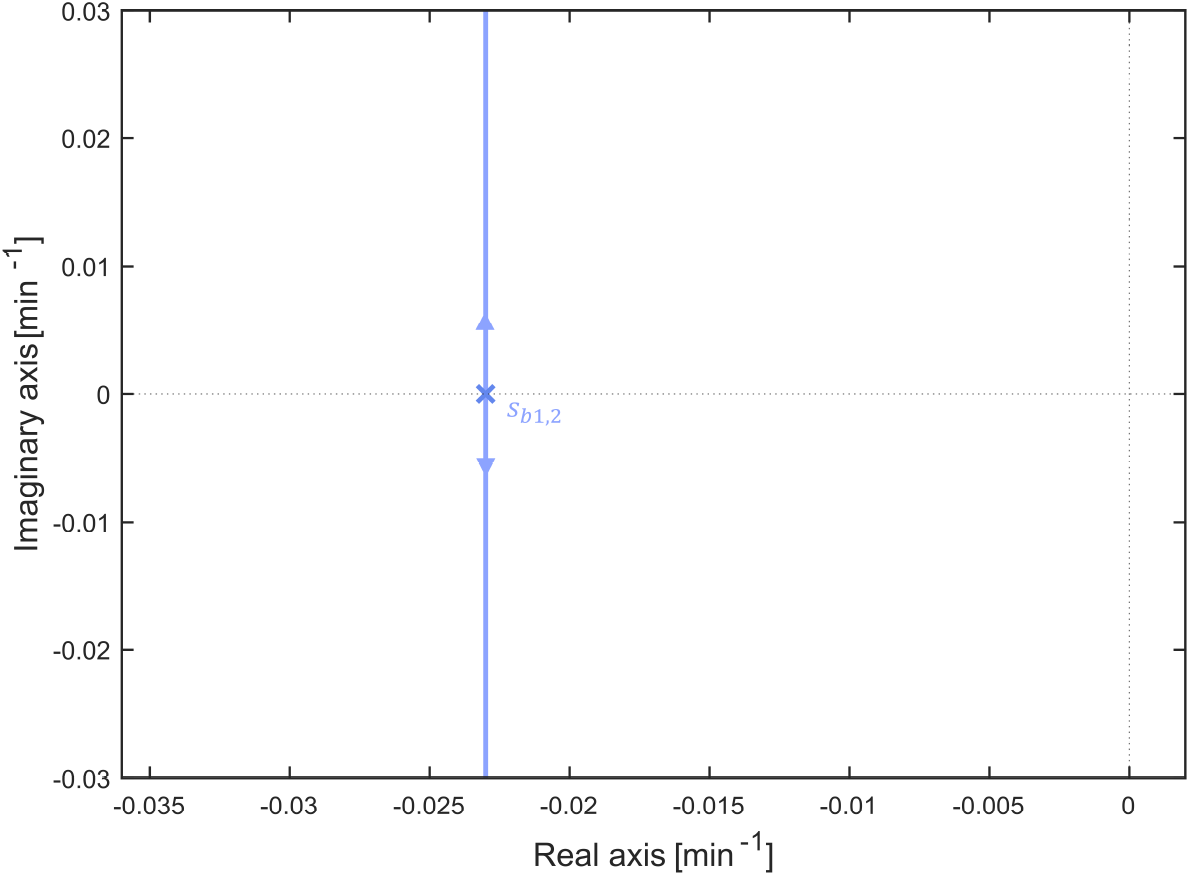
Root locus of the proposed multicellular P control architecture. Root locus of *k*_*P*_ *G*_*P*_ (*s*) (S65). The root locus has two branches starting from the poles and going to infinity. Different values of *k*_*P*_ select different closed-loop poles on the path.

